# Spatial regulation of coordinated excitatory and inhibitory synaptic plasticity at dendritic synapses

**DOI:** 10.1101/2021.05.31.446423

**Authors:** Tiziana Ravasenga, Massimo Ruben, Alice Polenghi, Enrica Maria Petrini, Andrea Barberis

## Abstract

The induction of synaptic plasticity at an individual dendritic glutamatergic spine can affect neighboring spines. This local modulation generates dendritic plasticity microdomains believed to expand the neuronal computational capacity. Here, we investigate whether local modulation of plasticity can also occur between glutamatergic synapses and adjacent GABAergic synapses. Using MNI-glutamate and DPNI-GABA double uncaging combined with electrophysiology, live-cell imaging and single-particle tracking, we find that the induction of LTP at an individual glutamatergic spine causes the depression of nearby GABAergic inhibitory synapses (within 3 microns), whereas more distant ones are potentiated. Notably, L-type calcium channels and calpain are required for this plasticity spreading. Overall, our data support a model whereby input-specific glutamatergic postsynaptic potentiation induces a spatially-regulated rearrangement of inhibitory synaptic strength in the surrounding area through short-range heterosynaptic interactions. Such local coordination of excitatory and inhibitory synaptic plasticity is expected to profoundly influence dendritic information processing and integration.

## Introduction

An increasing body of evidence shows that similarly to glutamatergic synapses, GABAergic synapses undergo many forms of short and long-term plasticity expressed at both the pre- and postsynaptic levels (Castillo et al., 2011; Chiu et al., 2018, 2019; Petrini and Barberis, 2014; Tyagarajan and Fritschy, 2014). This raises the important question of how excitatory and inhibitory synaptic plasticity are orchestrated during neuronal activity. Several studies show that the chronic modification of overall neuronal spiking activity can modulate both glutamatergic and GABAergic synapses within the same neuron, thus indicating the presence of cellular mechanisms coordinating activity-dependent changes in synaptic excitatory and inhibitory strength (Ibata et al., 2008; Rannals and Kapur, 2011; Turrigiano, 2011, 1999). However, the precise relationship between neuronal activity and the concomitant modifications of both glutamatergic and GABAergic synapses is poorly understood. This is mainly because the effects of diverse acute plasticity-inducing protocols have been mostly studied independently at either excitatory or inhibitory synapses.

Nevertheless, the induction and expression of glutamatergic and GABAergic plasticity show several common features. Postsynaptically, for instance, the activation of NMDA receptors and CaMKII, one of the best characterized signaling pathways in excitatory glutamatergic long-term potentiation (LTP) (Nicoll and Roche, 2013), is also crucial for the expression of inhibitory long-term potentiation (iLTP) (Chiu et al., 2018; Flores et al., 2015; He et al., 2015; Kano et al., 1996; Marsden et al., 2007; Petrini et al., 2014). Moreover, GABAergic postsynaptic depression relies on the activity of calcineurin (Bannai et al., 2009, 2015; Muir et al., 2010), a phosphatase that is also implicated in glutamatergic long-term depression (LTD) (Mulkey et al., 1994; Zeng et al., 2001). In addition the proteolytic action of calpains has been reported to affect both glutamatergic and GABAergic synaptic plasticity (Andres et al., 2013; Briz and Baudry, 2017; Costa et al., 2016; Tyagarajan et al., 2013). Along the same lines, postsynaptic GABAergic iLTP and iLTD involve the increase and decrease, respectively, of synaptic GABAA receptor number (Bannai et al., 2009; Chiu et al., 2018; He et al., 2015; Kurotani et al., 2008; Muir et al., 2010; Nusser et al., 1998; Petrini et al., 2014). Such a mechanism is analogous to that previously demonstrated at glutamatergic synapses, where the bidirectional tuning of postsynaptic AMPA receptor number positively and negatively modulates synaptic strength (Diering and Huganir, 2018). In addition, both glutamatergic and GABAergic synaptic plasticity depend on the regulated interaction of postsynaptic receptors with scaffold proteins which, in turn, modulate receptor lateral diffusion at synapses (Carta et al., 2013; Choquet and Triller, 2013; Petrini and Barberis, 2014).

Importantly, inhibitory postsynaptic plasticity occurs at GABAergic synapses located in dendrites of pyramidal neurons, i.e. at inhibitory synapses that, at least in specific dendritic sub-regions, can be located only few microns away from glutamatergic synapses (Chen et al., 2012; Chiu et al., 2018; Flores et al., 2015; Megías et al., 2001; Petrini et al., 2014; van Versendaal et al., 2012; Villa et al., 2016). Moreover, a subset of GABAergic synapses directly impinge onto glutamatergic spines, representing extremely close spatial localization between excitatory and inhibitory synapses (Chen et al., 2012; Chiu et al., 2013; Kubota et al., 2007; Nusser et al., 1996; Tamás et al., 2003; Villa et al., 2016). Thus, the analogous mechanisms of postsynaptic plasticity and the spatial contiguity of GABAergic and glutamatergic synapses suggest the hypothesis of local interplay in dendrites during synaptic plasticity between neighboring excitatory and inhibitory synapses. However, while it has been proposed that the plasticity-induced rearrangement of dendritic excitatory and inhibitory synapses can be clustered in specific dendritic subdomains (Chen et al., 2012), the nature of short-range heterosynaptic interactions between glutamatergic and GABAergic synapses remains to be elucidated.

In the present study, we investigated the spatial determinants involved in the interplay between plasticity at hippocampal dendritic excitatory and inhibitory synapses. We find that action potential trains at 2 Hz (low frequency stimulation, LFS) delivered to the postsynaptic neuron induce a non-associative inhibitory long-term synaptic potentiation (iLTP) which is concomitant with excitatory synaptic long-term depression (LTD). Interestingly, pairing LFS with MNI-glutamate uncaging at an individual glutamatergic spine (known to induce single-spine LTP) causes the long-term depression of GABAergic synapses (iLTD) located in the vicinity of the potentiated glutamatergic spine (within 3 μm), while more distant inhibitory synapses exhibit iLTP. The local iLTD – which spatially correlates with a confined dendritic calcium increase – can be selectively reverted to iLTP by the inhibition of L-type voltage gated calcium channels and of calpain activity, suggesting there is a functional plasticity microdomain near the potentiated spine. Notably, iLTD and iLTP at close and more distant GABAergic synapses show an opposite regulation of gephyrin clustering and synaptic GABAA receptor surface dynamics. These data demonstrate a high degree of coordination between plasticity at dendritic excitatory and inhibitory synapses and reveal that this functional interplay is restricted to dendritic microdomains.

## Results

### Electrophysiological induction of inhibitory long-term potentiation (iLTP)

We first sought to identify an electrical stimulation protocol to induce inhibitory long-term potentiation (iLTP) on cultured hippocampal neurons. To this end, using paired patch recordings, we tested different depolarization protocols on a pyramidal cell and monitored the amplitude changes of IPSCs elicited by an individual action potential in an identified parvalbumin-positive interneuron (PV+) (presynaptic neuron) connected with the stimulated pyramidal cell (postsynaptic neuron) (Figure 1A). Delivery of a 40 s action potential (AP) train at 2 Hz (low frequency stimulation, LFS) to pyramidal neurons induced a robust increase of inhibitory post-synaptic currents (IPSCs) amplitude (at 25-30 min: 1.27 ± 0.01 fold increase, n = 24, p<0.001; Figure 1A bottom and 1B) that lasted for more than 30 minutes. We next investigated whether this form of inhibitory long-term potentiation (iLTP) was expressed on the pre- or post-synaptic side. First, the amplitude ratio between two consecutive IPSCs elicited at a 50 ms interval (paired pulse ratio) was unchanged before and after the delivery of the LFS protocol (0.96 ± 0.02 and 0.94 ± 0.02, respectively, n = 25, p>0.05), suggesting a postsynaptic expression mechanism (Figure S1A). Next, we considered the possibility that, as described by previous studies (Guevara-Guzman et al., 2002; Lourenço et al., 2014; Saransaari and Oja, 2006; Xue et al., 2011), the potentiation of IPSCs could rely on the increased GABA release induced by the presynaptic action of nitric oxide (NO). The application of the nitric oxide synthase blocker L-NAME was unable to prevent the LFS-induced iLTP (at 25-30 min: normalized IPSCs amplitude = 1.40 ± 0.02 of baseline, n = 5, p=0.03; Figure S1B), thus making a presynaptic mechanism for this form of GABAergic synaptic potentiation unlikely.

**Figure 1:**
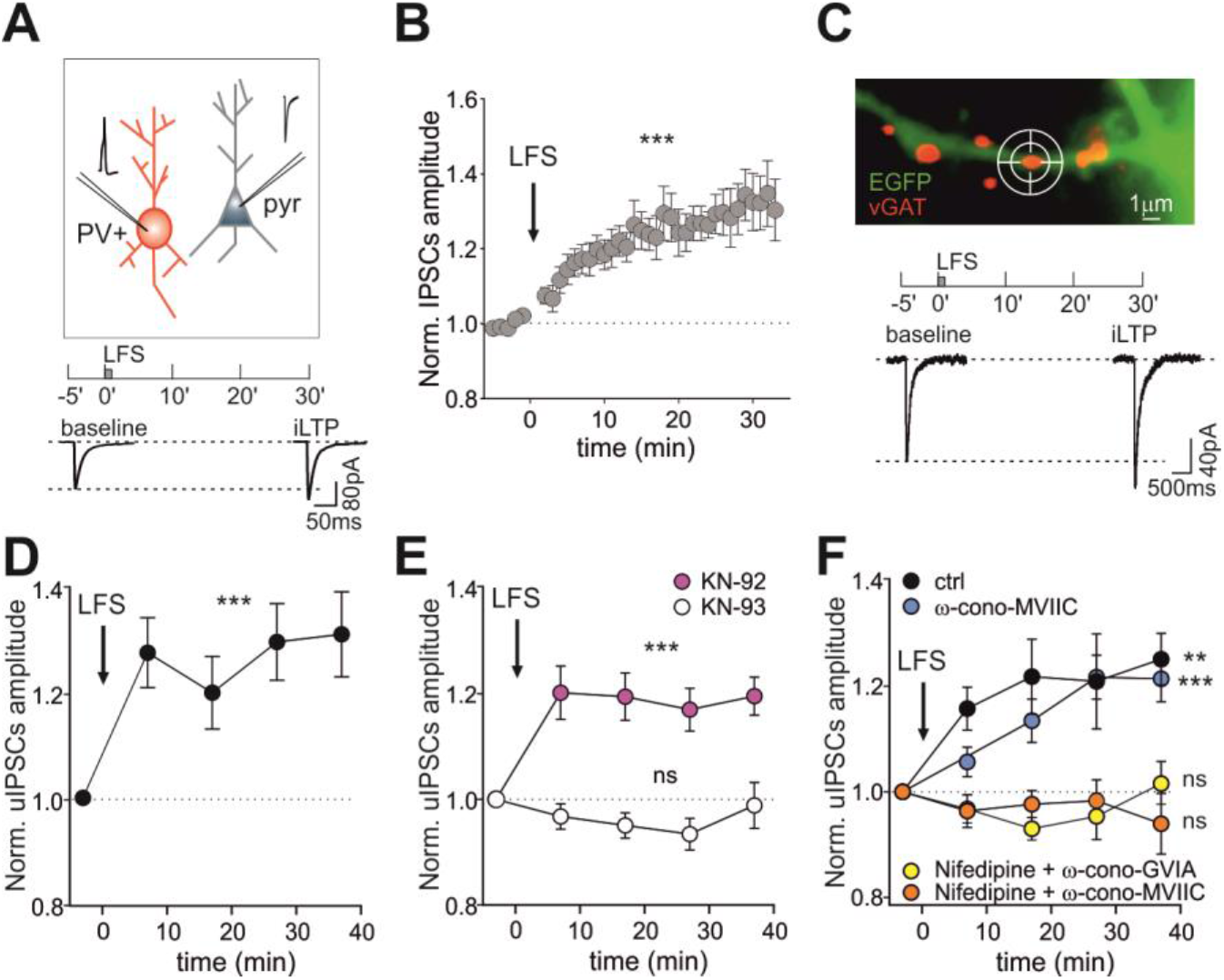
LFS induces iLTP. **A.** Top: Experimental configuration of paired patch recordings including a presynaptic parvalbumin-tdTomato positive (PV+) interneuron (red) and a postsynaptic pyramidal cell (grey). The schematic shows the low-frequency protocol (2 Hz APs train for 40 s, LFS) to induce synaptic plasticity. Bottom: Representative average traces of inhibitory postsynaptic response (IPSCs) before and 30 min after the protocol. **B.** Potentiation of IPSC amplitude after LFS (arrow; n = 24 neurons, F_37,708_ = 5.3, p < 0.001, one-way ANOVA followed by Turkey’s multiple comparison test post-hoc test). **C.** Top: Identification of GABAergic synapses by live labeling of vGAT (red, see Star Methods) in an EGFP-expressing neuron (green). The “target” symbol indicates an individual GABAergic synapse where a diffraction-limited 378 nm UV laser spot was directed to uncage DPNI-GABA (see Star Methods). Scale bar, 1 μm. Timeline of the experiment (LFS, as in A). Bottom: Representative averaged traces of uncaging IPSCs (uIPSCs) before (baseline) and 30 min after LFS (iLTP). **D.** uIPSCs are potentiated after LFS (n = 23 synapses from 7 neurons; F_4,87_ = 5.0, p = 0.001; one-way ANOVA followed by Dunnett’s post-test). **E.** CaMKII is required for LFS-induced iLTP. uIPSC amplitude normalized to baseline values in the presence of KN-93 (white; n = 19 synapses from 5 neurons; F_4,61_ = 1.4, p = 0.24, one-way ANOVA followed by Dunnett’s post-test) and the inactive analogue KN-92 (pink; n = 26 synapses from 7 neurons; F_4,92_ = 6.5, p < 0.001; one-way ANOVA followed by Dunnett’s post-test). **F.** Influence of voltage-gated calcium channels (VGCCs) on iLTP expression. Relative (after/before) uIPSC amplitude upon LFS in control conditions (black; n = 24 synapses from 9 neurons; F_4,78_ = 5.1, p = 0.001), or in the presence of the following VGCCs blockers: ω-conotoxin MVIIC for P/Q and N-type (blue; n =16 synapses from 4 neurons; F_4,64_ = 9.1, p < 0.001), or nifedipine, for L-type and ω-conotoxin MVIIC (orange; n = 24 synapses from 6 neurons; F_4,80_ = 0.3, p = 0.88) or nifedipine and ω-conotoxin GVIA for N-type (yellow; n = 17 synapses from 6 neurons; F_4,52_ = 1.9, p = 0.13). All statistical comparisons were performed with one-way ANOVA followed by Dunnett’s post-test. Values are expressed as mean ± SEM. **p < 0.01, ***p < 0.001, ns = not significant. See also Figure S1.

Next, we found that the inclusion of the calcium chelator BAPTA in the patch pipette prevented the IPSC amplitude increase (0.97 ± 0.04 fold of baseline, n = 4, p>0.05; Figure S1C), indicating that this iLTP depends on the elevation of intracellular calcium following delivery of the LFS. However, blocking NMDA receptors or L-type voltage-gated calcium channels (VGCCs) by APV or nifedipine, respectively, did not prevent the LFS-dependent increase in GABAergic currents (APV: 1.16 ± 0.01 fold increase of baseline, n = 11, p<0.001; Figure S1D and nifedipine: 1.16 ± 0.01 fold of baseline, n = 21, p<0.001; Figure S1E). Likewise, co-application of APV and nifedipine failed to inhibit iLTP (normalized IPSC amplitude = 1.13 ± 0.03 of baseline, n = 6, p=0.002; Figure S1F). Worth noting, additional potential sources of calcium influx mediated by N-, P/Q- and R-type VGCC (also expressed presynaptically) were not investigated in these paired-patch configuration experiments, since their blockade could affect GABA release and the amplitude of IPSCs, thus masking their role in iLTP induction.

We corroborated the postsynaptic expression of this LFS-induced iLTP with an optical approach based on the photolysis of caged DPNI-GABA by UV light focused in diffraction-limited spots. With this technique, it is possible to exogenously elicit uncaging IPSCs (uIPSCs) at GABAergic synapses with single-synapse specificity and to keep constant the trial-to-trial amount of GABA delivered to the postsynaptic element (see STAR methods). In the present experiments, inhibitory synapses were identified by live immunolabeling of the presynaptic marker vesicular GABA transporter (vGAT) (see STAR methods) (Figure 1C, top). Consistently with the paired-patch experiments, after the application of the LFS protocol, the amplitude of uIPSCs elicited by DPNI-GABA uncaging at dendritic proximal inhibitory synapses was significantly increased (at 27 min: 1.30 ± 0.07 fold increase of baseline, n = 23 from 7 neurons, p=0.001; Figure 1C bottom and 1D). Overall, these data demonstrate that GABAergic long-term potentiation induced by LFS is expressed on the postsynaptic side.

We then investigated whether, similarly to previous studies (Chiu et al., 2018; Flores et al., 2015; Marsden et al., 2007; Petrini et al., 2014), the activation of CaMKII could be involved in the expression of this form of iLTP. We found that the LFS-induced iLTP was prevented by the application of the CaMKII inhibitor KN-93 (normalized uIPSC amplitude at 27 min: 0.93 ± 0.03, n = 19 from 5 neurons, p>0.05), whereas KN-92, an inactive analogue of KN-93, left iLTP expression unchanged (normalized uIPSC amplitude at 27 min: 1.17 ± 0.04, n = 26 from 7 neurons, p<0.001, Figure 1E). Since as stated above, N-, P/Q- and R-type VGCCs can potentially be located pre- or post-synaptically, we further investigated whether this iLTP depends on dendritic calcium elevation mediated by the postsynaptic N-, P/Q- and R-type VGCCs by taking advantage of the uncaging technique, which induces synaptic-like currents without engaging the presynaptic release machinery. While in the presence of ω-conotoxin MVIIC to block the P/Q- and N-type VGCCs we observed iLTP similarly to control conditions without drugs (uIPSCs amplitude at 27 min: ctrl= 1.21 ± 0.09 fold increase, n = 24 from 9 neurons, p=0.001; MVIIC= 1.22 ± 0.04 fold increase, n = 16 from 4 neurons, p<0.001), the co-application of nifedipine and ω-conotoxin MVIIC to block L-along with P/Q- and N-type VGCCs abolished it (0.97 ± 0.04 fold increase, n = 24 from 6 neurons, p>0.05; Figure 1F). Further dissecting the calcium sources responsible for iLTP induction, ω- conotoxin GVIA, a specific blocker for N-type VGCCs, combined with the L-type VGCCs nifedipine prevented iLTP (normalized uIPSC amplitude at 27 min: 0.95 ± 0.04 fold increase, n = 17 from 6 neurons, p>0.05; Figure 1F), indicating the concomitant involvement of L- and N-type VGCC with a negligible role for P/Q-type VGCC. This finding is in line with previous reports showing that P/Q-type VGCC are poorly expressed in dendrites of hippocampal pyramidal neurons (Higley and Sabatini, 2008). Overall, these results suggest that the application of LFS promotes the increase of IPSC amplitude via the increase of intracellular calcium concentration through the L- and N-type VGCC and the activation of CaMKII.

### LFS induces long-term depression at glutamatergic synapses (LTD)

Next, we studied the plasticity of glutamatergic synapses in response to the same LFS protocol used to induce GABAergic iLTP. To this end, in a paired-patch configuration between two connected pyramidal neurons, we elicited a single AP in the presynaptic neuron and recorded the consequent excitatory post-synaptic current (EPSC) in the postsynaptic neuron (Figure 2A). The delivery of the LFS protocol to the postsynaptic neuron was responsible for a persistent and significant reduction of EPSC amplitude (at 25-30 min: 0.69 ± 0.03 fold of baseline, n = 15, p<001), thus indicating the expression of glutamatergic long-term depression (LTD) (Figure 2B). To assess whether this plasticity of glutamatergic synapses was expressed at the postsynaptic level, we photo-released MNI-glutamate at individual glutamatergic synapses at spines identified by overexpressed Homer1c-GFP, a GFP-tagged component of the glutamatergic postsynaptic density (PSD) (Fig. 2C, top). We found that after LFS, the amplitude of uncaging EPSCs (uEPSCs) was significantly reduced (at 27 min: 0.82 ± 0.04 fold of baseline, n = 16 from 11 neurons, p<0.001), thus indicating that this protocol induced a postsynaptic form of glutamatergic LTD (Figure 2C, bottom, Figure 2D). This observation is in line with previous studies demonstrating that low frequency stimulations (in the range of 0.5-5 Hz) are responsible for glutamatergic LTD (Dudek and Bear, 1992; Malenka and Bear, 2004). Overall, we found that the delivery of LFS protocols was sufficient to induce a non-Hebbian potentiation of GABAergic (iLTP) and the depression of glutamatergic synapses (LTD), thus leading to an opposite regulation of the plasticity of excitatory and inhibitory synapses.

**Figure 2:**
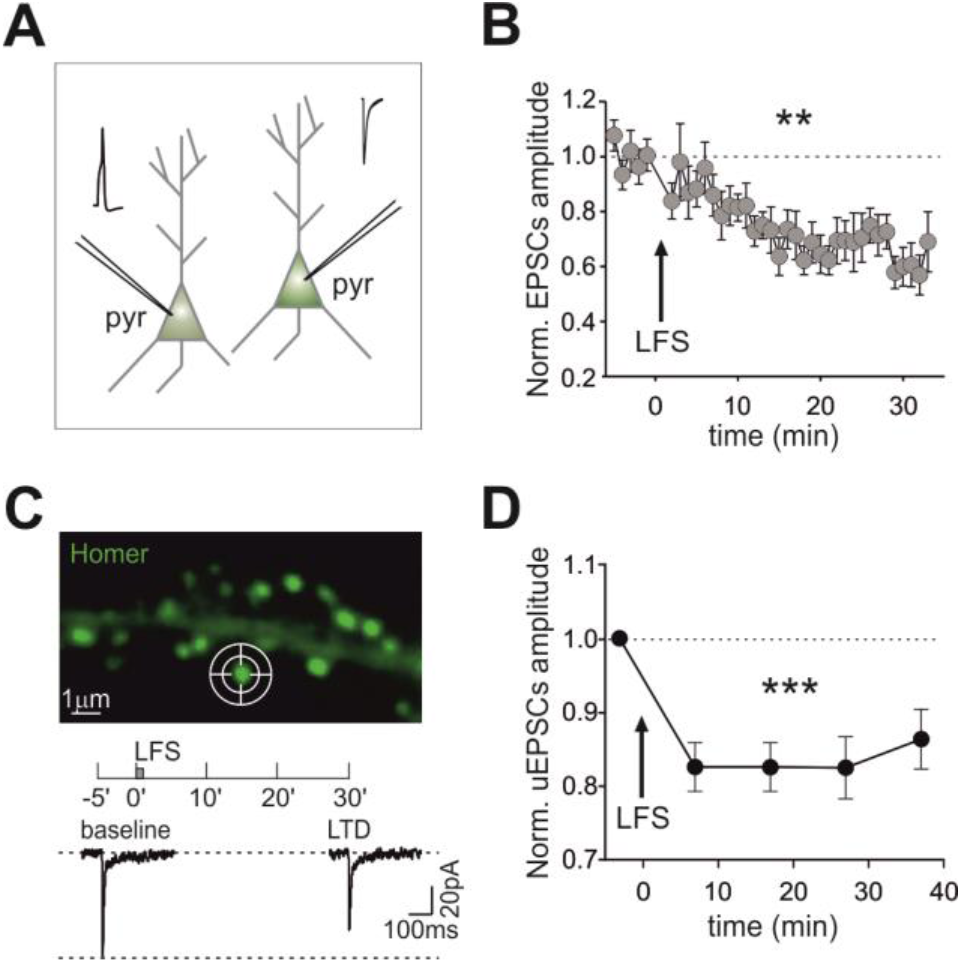
iLTP-inducing protocol promotes LTD at excitatory synapses. **A.** Experimental configuration of paired patch recordings used to probe excitatory postsynaptic currents (EPSCs) in a pyramidal neuron interconnected with another pyramidal cell. Synaptic plasticity was induced following the protocol outlined in Figure 1A. **B.** LFS induces long-term depression (LTD). Time course of normalized EPSC amplitude after LFS delivery (arrow) over 30 min (n = 15 synapses, F_39,438_ = 2.8, p < 0.001, one-way ANOVA followed by Turkey’s test). **C.** Top: Identification of glutamatergic spines by Homer1c-GFP fluorescence. Scale bar, 1 μm. The target symbol indicates where a diffraction-limited 378 nm UV laser spot was directed to uncage MNI-glutamate at an individual spine. Bottom: Timeline of the experiment (as in Figure 1C) and representative average uncaging EPSC (uEPSC) traces recorded before (baseline) and at 30 minutes after LFS (LTD). **D.** Persistent reduction of uEPSC amplitude upon LFS as compared to baseline (n = 16 synapses from 11 neurons, F_4,68_ = 6.0, p < 0.001; one-way ANOVA followed by Dunnett’s post-test). Values are expressed as mean ± SEM. **p < 0.01, ***p < 0.001.

### Increase of gephyrin clusters is associated with iLTP expression

We previously demonstrated that the postsynaptic accumulation of the scaffold protein gephyrin is involved in the expression of a chemically-induced postsynaptic potentiation of GABAergic currents (chem-iLTP) (Petrini et al., 2014). We therefore investigated whether the expression of this electrically-induced iLTP similarly involves the rearrangement of synaptic gephyrin. To this end, we monitored gephyrin-GFP fluorescence over time at individual clusters before and after the application of the iLTP induction protocol (Figure 3A and 3B). After the LFS, the fluorescence intensity of individual gephyrin-GFP clusters significantly increased with respect to the baseline values (1.17 ± 0.03 fold increase, n = 13, p<0.001; Figure 3B and 3C), reflecting a promoted accumulation of synaptic gephyrin during iLTP. Collectively, the potentiated GABAergic synaptic currents concomitant with increased synaptic gephyrin upon LFS indicates that, similarly to previous studies, postsynaptically expressed iLTP may involve the enrichment of the inhibitory PSD (iPSD).

**Figure 3:**
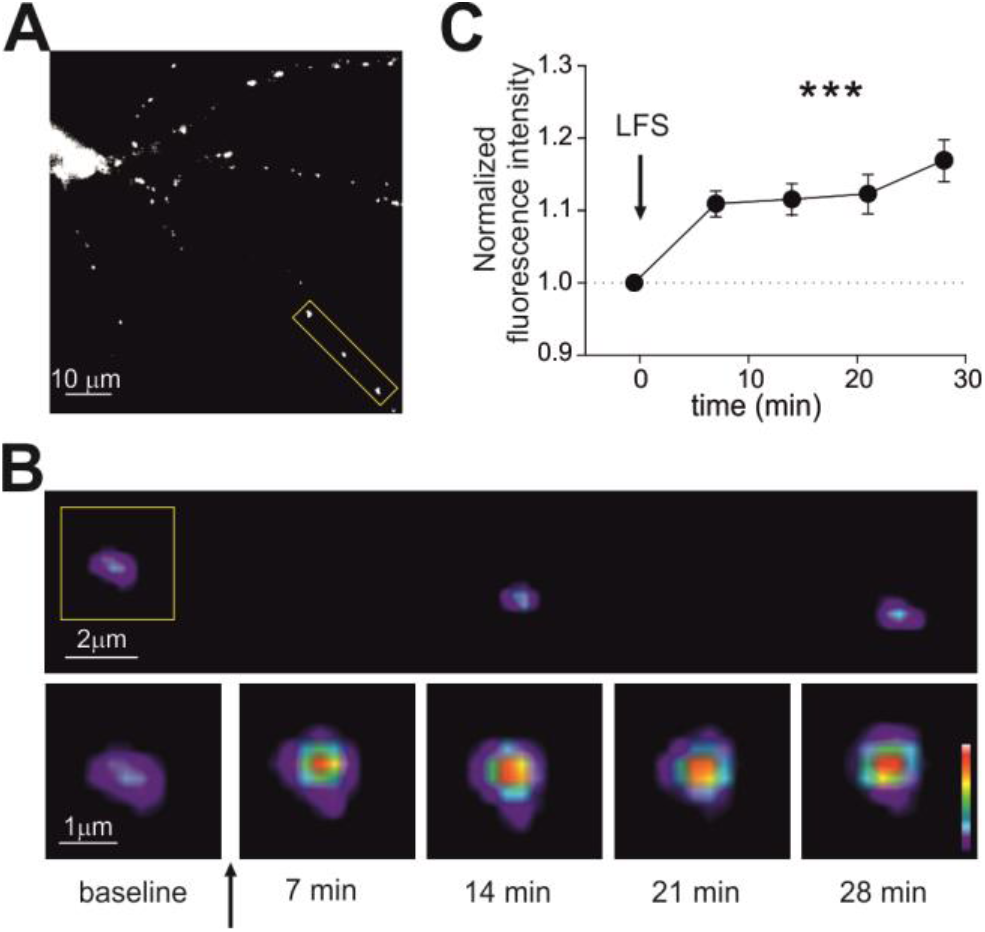
Enhanced gephyrin clustering during iLTP. **A.** Representative epifluorescence image of a neuron expressing GFP-tagged gephyrin. Scale bar, 10 μm. **B.** Top: Pseudocolor magnification of the dendritic portion framed in A. Scale bar, 2 μm. Please note that the fluorescence scale has been enhanced to visualize small clusters. Bottom: Pseudocolor images of the gephyrin cluster framed above at different time points before and after LFS (arrow). Scale bar, 1 μm. **C.** Summary of the normalized gephyrin fluorescence increase (after/before) observed upon iLTP induction with LFS (arrow; n = 13, F_4,48_ = 21.5,, p < 0.001, RM one-way ANOVA followed by Dunnett’s post-test). Values are expressed as mean ± SEM. ***p < 0.001.

### Interaction between plasticity at excitatory and inhibitory synapses

We next analyzed the effect of the expression of glutamatergic excitatory long-term potentiation (LTP) on GABAergic synaptic function, in particular testing if the distance between dendritic excitatory and inhibitory synapses could be a determinant for their activity-dependent interplay. We first induced “single-spine LTP” by pairing the depolarization of a pyramidal cell (obtained with LFS) with repetitive uncaging of MNI-glutamate at an individual glutamatergic spine (see STAR methods). As such, we could mimic a Hebbian stimulation paradigm where the postsynaptic depolarization is concomitant with the presynaptic glutamate release at a specific input (Harvey and Svoboda, 2007; Matsuzaki et al., 2004). Next, in a double uncaging experiment we photo-released both MNI-glutamate and DPNI-GABA to probe the strength of glutamatergic and GABAergic synapses, respectively upon the induction of single-spine LTP. In a typical experimental layout (Figure 4A, top), we considered two glutamatergic and two GABAergic synapses identified by the overexpression of Homer1c-DsRed and gephyrin-GFP, respectively, chosen at different relative distances on the same dendrite. We found that the delivery of the LFS protocol along with single spine MNI-glutamate laser uncaging at 4 Hz (Figure 4A, bottom) selectively increased the amplitude of uEPSCs elicited at the photo-stimulated spine (Figure 4B, top left, synapse #1), thus indicating the expression of single-spine LTP (uEPSC amplitude: 1.20 ± 0.05 fold increase, n = 7- 20 synapses from 20 neurons, p<0.001; Figure S2A). Overall, after the delivery of the single-spine LTP protocol, ∼ 84% of the stimulated synapses exhibited long-term potentiation, whereas glutamatergic synapses located on spines other than the photo-stimulated one showed LTD (Figure 4B, top right, synapse #3; normalized uEPSC amplitude: 0.88 ± 0.04 of baseline, n = 6-16 synapses from 20 neurons, p<0.001; Figure S2A). Concomitantly, GABAergic synapses located at distance > 3 μm from the photo-stimulated spine were potentiated (iLTP) (Figure 4B, bottom right, synapse #4; normalized uIPSC amplitude: 1.21 ± 0.05 of baseline, n = 7-41 synapses from 20 neurons, p<0.001; Figure S2B). This plasticity pattern (i.e. LTD and concomitant iLTP) closely matches that obtained at GABAergic and glutamatergic synapses by the non-Hebbian delivery of LFS shown in Figure 1 and Figure 2. Intriguingly, GABAergic synapses located in the close vicinity of the potentiated spine (< 3 μm) were depressed (iLTD) (Figure 4B bottom left, synapse #2; normalized uIPSC amplitude: 0.89 ± 0.04 of baseline, n = 11-30 synapses from 20 neurons, p=0.02; Figure S2B), thus showing that inhibitory synaptic plasticity significantly reverts its sign at increasing distance from the potentiated spine (Figure 4C, green, for each data-point n = 18-39 from 20 neurons, p<0.001). This suggests that the long-term potentiation of an individual glutamatergic spine generates a region of reduced inhibition in the spatial range of ± 3μm.

**Figure 4:**
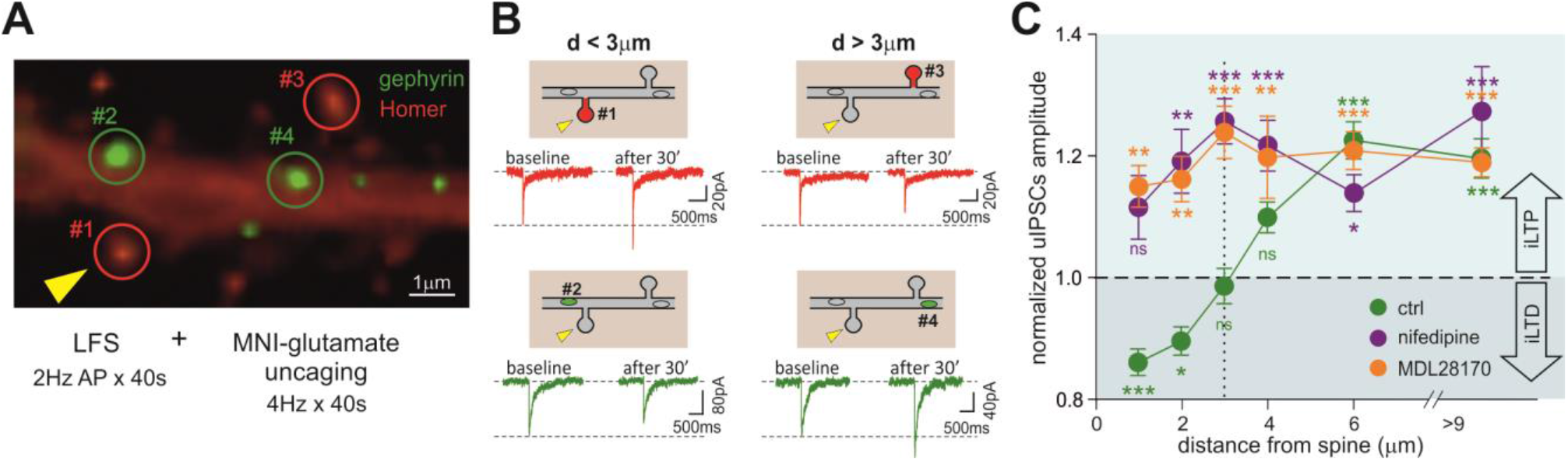
Plasticity interplay between potentiated spine and neighboring GABAergic synapses. **A.** Epifluorescence image showing a typical experimental layout. Top: Dendritic portion of a neuron expressing Homer1c-DsRed (red) to identify excitatory spines and Gephyrin-GFP (green) to identify inhibitory synapses. Scale bar, 1 μm. Excitatory and inhibitory synaptic responses were probed by uncaging MNI-glutamate at two glutamatergic spines (#1 and #3) and DPNI-GABA at two GABAergic synapses (#2 and #4), respectively. The yellow arrowhead indicates the stimulated spine (see below). Bottom: Induction of synaptic plasticity. LFS (2Hz AP train) delivered to the whole neuron through the patch pipette was paired with diffraction-limited 4 Hz MNI-glutamate uncaging (yellow arrowhead) selectively at spine #1 (LFS + MNI-glutamate uncaging). **B.** Representative average traces of uEPSCs (top, red) and uIPSCs (bottom, green) recorded from glutamatergic synapses (#1 and #3) and GABAergic synapses (#2 and #4) before (baseline) and 30 min after the induction of synaptic plasticity. The relative localization of each spine with respect to #1 (receiving the single spine LTP protocol, yellow arrowhead) is schematized above the traces. Please note that the stimulated synapse #1 displays LTP, while glutamatergic synapse #3 and GABAergic synapse #4, both located relatively far from the potentiated spine, show LTD and iLTP, respectively. Interestingly, GABAergic synapse #2, close to the potentiated spine, shows iLTD. **C.** Spatial distribution of GABAergic plasticity at inhibitory synapses located at different distances from the potentiated spine. Please note that uIPSCs at GABAergic synapses located in close proximity of the stimulated spine (d < 3 μm) were depressed, whereas those located at d > 3 μm displayed potentiation (green, for each data point n = 18-39 synapses from 20 neurons, F_6,603_ = 30.6, p< 0.001, one-way ANOVA followed by Dunnett’s post-test). In the presence of nifedipine, the same protocol (LFS + MNI-glutamate uncaging) does not elicit iLTD at synapses at d < 3 μm. In these conditions, all GABAergic synapses exhibit iLTP regardless of their distance from the stimulated spine (purple, for each data point n = 7-16 synapses from 7 neurons, F_6,68_ = 6.6, p < 0.001, one-way ANOVA followed by Dunnett’s post-test). The blockade of calpain activity with MDL28170 prevents the local iLTD (orange, for each data point n = 9-67 synapses from 24 neurons, F_6,223_ = 13.0, p < 0.001, one-way ANOVA followed by Dunnett’s post-test). Values are expressed as mean ± SEM.*p < 0.05, **p < 0.01, ***p < 0.001, ns = not significant. See also Figure S2.

Next, we studied the role of calcium in this plasticity interplay observed at excitatory and inhibitory synapses upon the delivery of the “LFS + MNI-glutamate uncaging” to induce single-spine LTP. First, we explored the contribution of L-type VGCCs. After assessing that nifedipine did not affect the expression of single-spine LTP (uEPSC amplitude: 1.32 ± 0.08 fold increase, n = 4-7 from 7 neurons, p = 0.01; Figure S2C), we observed that the blockade of L-type VGCC prevented local GABAergic iLTD at distances < 3 μm from the potentiated spine and reverted it to iLTP, while leaving the iLTP at distance > 3 μm unaffected (Figure 4C, purple, for each data-point n = 7-16 synapses from 7 neurons). More specifically, at 27 minutes, uIPSCs amplitude at synapses close to the potentiated spine was 1.22 ± 0.03 of baseline (n = 3-9 from 7 neurons, p=0.01; Figure S2D), and at more distant GABAergic synapses was 1.16 ± 0.04 of baseline, (n = 4-14 from 7 neurons, p<0.001; Figure S2D). Overall, we demonstrate that while the application of LFS elicits diffuse inhibitory long-term potentiation, the pairing of LFS with MNI-glutamate uncaging at individual spines induces single-spine LTP and determines a local iLTD restricted to a range of 3 μm from the potentiated glutamatergic spine. Notably, this local iLTD as well as the depression of glutamatergic synapses depend on calcium entry through L-type VGCC.

### Calcium dynamics during the induction of excitatory and inhibitory plasticity

Next, we studied the spatial and temporal profile of dendritic calcium transients induced in the same neuron by either LFS protocols (non-Hebbian stimulation) or LFS paired with MNI-glutamate uncaging to induce single-spine LTP (Hebbian stimulation), which cause the two forms of plasticity interplay between glutamatergic and GABAergic synapses shown above (see STAR methods). We filled pyramidal cells with the synthetic calcium indicator Rhod-2 through the patch pipette and measured the dynamics of dendritic calcium during the delivery of these two plasticity-inducing protocols (Figure 5A). In response to 10 s of the LFS protocol, calcium rapidly increased and reached a plateau after approximately 3-5 s (Figure 5B black). In these conditions, the calcium increase was rather uniform along the dendrites (Figure 5C and 5D, top). The pairing of LFS with MNI-glutamate uncaging at an individual glutamatergic spine (identified by Homer1c-GFP fluorescence) (Figure 5A) induced an overall calcium increase that was considerably higher than that obtained by LFS stimulation alone, reaching a plateau after ∼ 5s (Figure 5B) and peaking in the dendritic portion below the photo-stimulated spine (Figure 5C and 5D, bottom). Next, we quantified the relative calcium increase induced by LFS+MNI-glutamate uncaging with respect to that induced by LFS in dendritic regions of 14 μm centered at the photo-stimulated spine (see STAR methods). Dendritic calcium concentration was significantly increased right below the spine (relative fluorescence intensity = 1.6 ± 0.15, n = 22, p<0.01; Figure 5E) and declined back to the calcium level induced by LFS within few micrometers. Importantly, such a range of confined dendritic calcium increase was on par with the spread of the local iLTD (Figure 4C) upon the induction of single-spine LTP. This finding supports the hypothesis that the spatially limited calcium increase in the vicinity of a potentiated glutamatergic spine could determine the confined iLTD expression.

**Figure 5:**
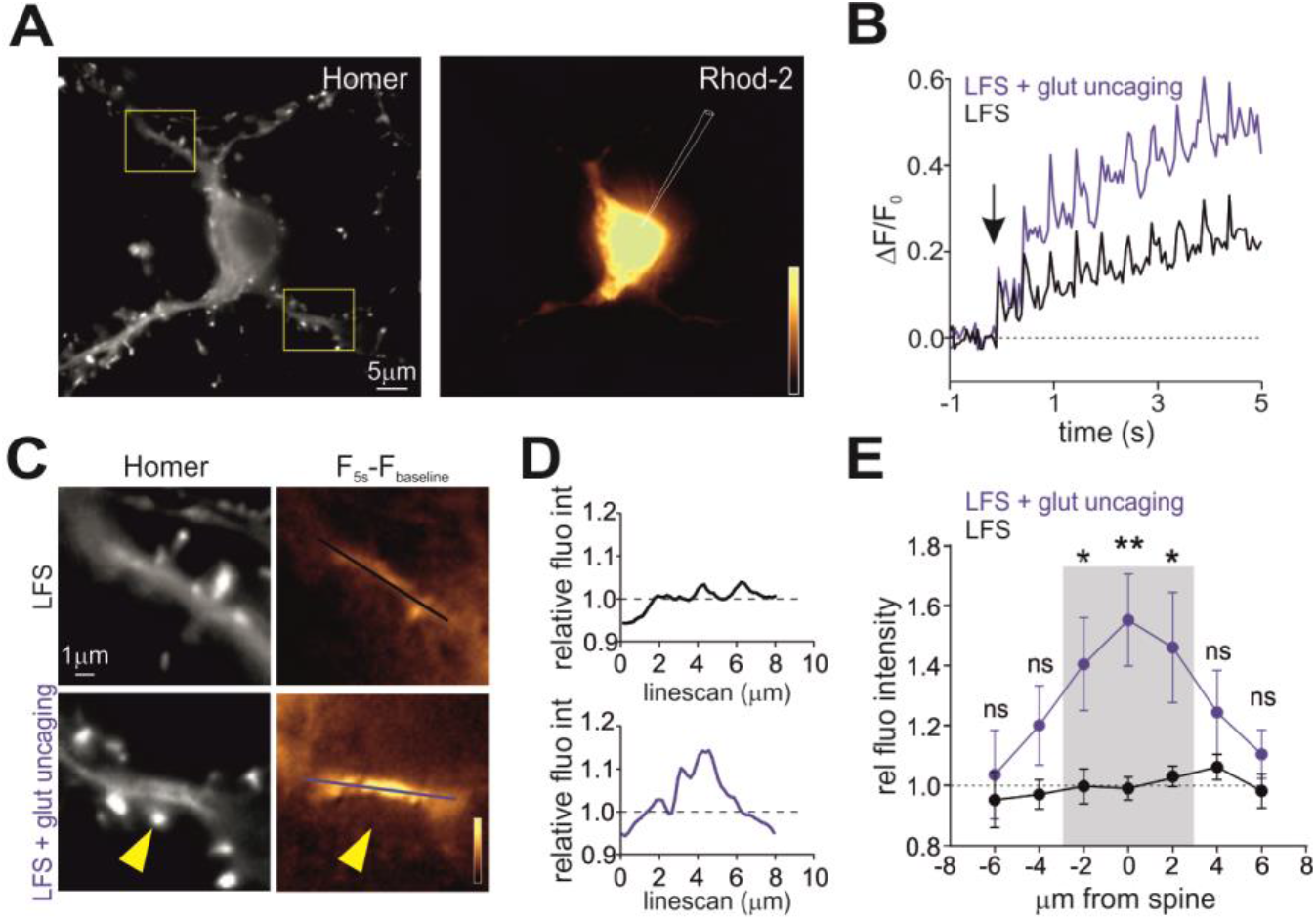
Spatial dynamics of dendritic calcium during iLTP and LTD. **A.** Representative epifluorescence image of a Homer1c-GFP expressing neuron (left) loaded with Rhod-2 through the patch pipette (gold, right). Scale bar, 5 μm. **B.** Relative Rhod-2 fluorescence intensity quantified during the LFS protocol (black) and the LFS paired with glutamate uncaging protocol (blue) in two 4 μm-long dendritic portions of the same neuron centered below a reference and stimulated spine, respectively. The arrow indicates the beginning of the protocol. **C.** Left: Magnifications of the dendritic portions framed in A, stimulated with LFS (top) or LFS paired with MNI-glutamate uncaging (bottom). The yellow arrowhead indicates the stimulated spine. Scale bar, 1 μm. Right: Gold pseudocolor representation of Rhod-2 fluorescence intensity changes at plateau (5 s) of the stimulating protocols (i.e., LFS, top and LFS paired with glutamate uncaging, bottom) with respect to baseline values (F_5s_-F_baseline_). The lines indicate the position of the linescans quantified in D. **D**. Relative fluorescence variation induced by “LFS + glut uncaging” protocol with respect to LFS alone. The fluorescence intensities quantified along the two linescans in C are normalized to the average fluorescence detected along the linescan in LFS. **E.** Changes in the relative dendritic Rhod-2 fluorescence intensity (as measured in Figure 5D) as a function of the distance from a reference or stimulated spine during the LFS (black) or the LFS+ glut uncaging (blue), respectively. The grey area indicates the range of ± 3μm from the potentiated spine where significant changes in Rhod-2 fluorescence are quantified as compared to the LFS protocol (LFS: n = 23 neurons, LFS+ glut uncaging: n = 22 neurons, F_1,265_ = 22.1, p < 0.001, two-way ANOVA followed by Bonferroni’s multiple comparison test). Statistical significance for each data point is shown. Values are expressed as mean ± SEM. *p < 0.05, **p < 0.01, ns = not significant.

In line with this hypothesis, we explored the possibility that calpains, a family of calcium-dependent proteases, could be involved in local iLTD. To test this possibility, we interfered with calpain activity by treating neurons with the broad range calpain inhibitor III (MDL 28170) before delivering the LFS + glutamate uncaging protocol to induce single-spine LTP. Under these conditions, we studied synaptic plasticity of inhibitory synapses while considering their distance from the potentiated spine (Figure 4C, orange, for each data-point n = 9-67 synapses from 24 neurons). GABAergic synapses close to the potentiated spine (d < 3 μm) not only did they lack the depression (local iLTD) observed in control conditions (Figure 4C, compare orange and green), but they in fact exhibited iLTP (d < 3μm: uIPSC amplitude = 1.19 ± 0.08 of baseline, n = 5-24 synapses from 24 neurons, p<0.001; Figure S2F). Interestingly, upon calpain blockade, the temporal profiles of iLTP expression at inhibitory synapses located at d < 3 μm and at d > 3 μm from the potentiated spine were comparable (Figure 4C orange) (d > 3μm: uIPSC amplitude =1.24 ± 0.04 of baseline, n = 6-68 synapses from 24 neurons, p<0.001; Figure S2F), appearing similar to neurons only exposed to LFS (compare with Figure 1). This evidence suggests that calpain blockade interferes with the confined effects of MNI-glutamate uncaging on inhibitory synaptic plasticity.

### Gephyrin dynamics during the expression of single-spine LTP

Considering that the expression of LFS-induced iLTP involves the promoted clustering of gephyrin at inhibitory synapses (Figure 3), we next investigated whether the converse was true, that is, if the iLTD observed nearby a potentiated glutamatergic spine is associated with the loss of synaptic gephyrin. We identified gephyrin clusters by means of FingRs (Fibronectin intrabodies generated with mRNA display) intrabodies (Gross et al., 2013) fused with GFP to label endogenous gephyrin. While still using GFP signal as a proxy for gephyrin, the intrabodies-based staining procedure significantly reduced gephyrin background fluorescence as compared to gephyrin-GFP overexpression, leading to a higher signal-to-noise ratio that proved essential to satisfactorily detect the mild gephyrin fluorescence reductions associated with iLTD. Similarly to the experiments shown in Figure 4, we induced single-spine LTP by pairing the LFS protocol with repetitive MNI-uncaging at an individual spine (identified by Homer1c-DsRed overexpression) and quantified GFP fluorescence over time at gephyrin clusters either close (d < 3 μm) or far (d > 3 μm) from the potentiated spine (Figure 6A). We found that after the delivery of the paired LFS and MNI-glutamate uncaging protocols, gephyrin clusters at d < 3 μm from the potentiated spine were significantly reduced (gephyrin normalized fluorescence intensity = 0.90 ± 0.04 of baseline, n = 13, p=0.002; Figure 6B), whereas at further distances they were increased (gephyrin normalized fluorescence intensity = 1.08 ± 0.04 of baseline, n = 13, p=0.04; Figure 6C). These results indicate that, following the delivery of the Hebbian-like stimulations, the spatial segregation of positive and negative plastic changes of GABAergic uIPSC amplitude closely matches changes in gephyrin accumulation. This indicates that the regulation of local gephyrin clustering could underlie the spatial dependence of GABAergic synaptic plasticity. The extent of potentiation of gephyrin clusters upon LFS is similar to that observed at distances > 3 μm from the potentiated spine upon the delivery of the LFS + MNI-glutamate uncaging paired protocol. This suggests that the increase of gephyrin fluorescence associated with iLTP can be similarly detected using gephyrin-GFP overexpression or the gephyrin FingRs-GFP strategy (compare Figure 3C and Figure 6C). This observation corroborates the match between electrophysiological, calcium imaging and gephyrin imaging data.

**Figure 6:**
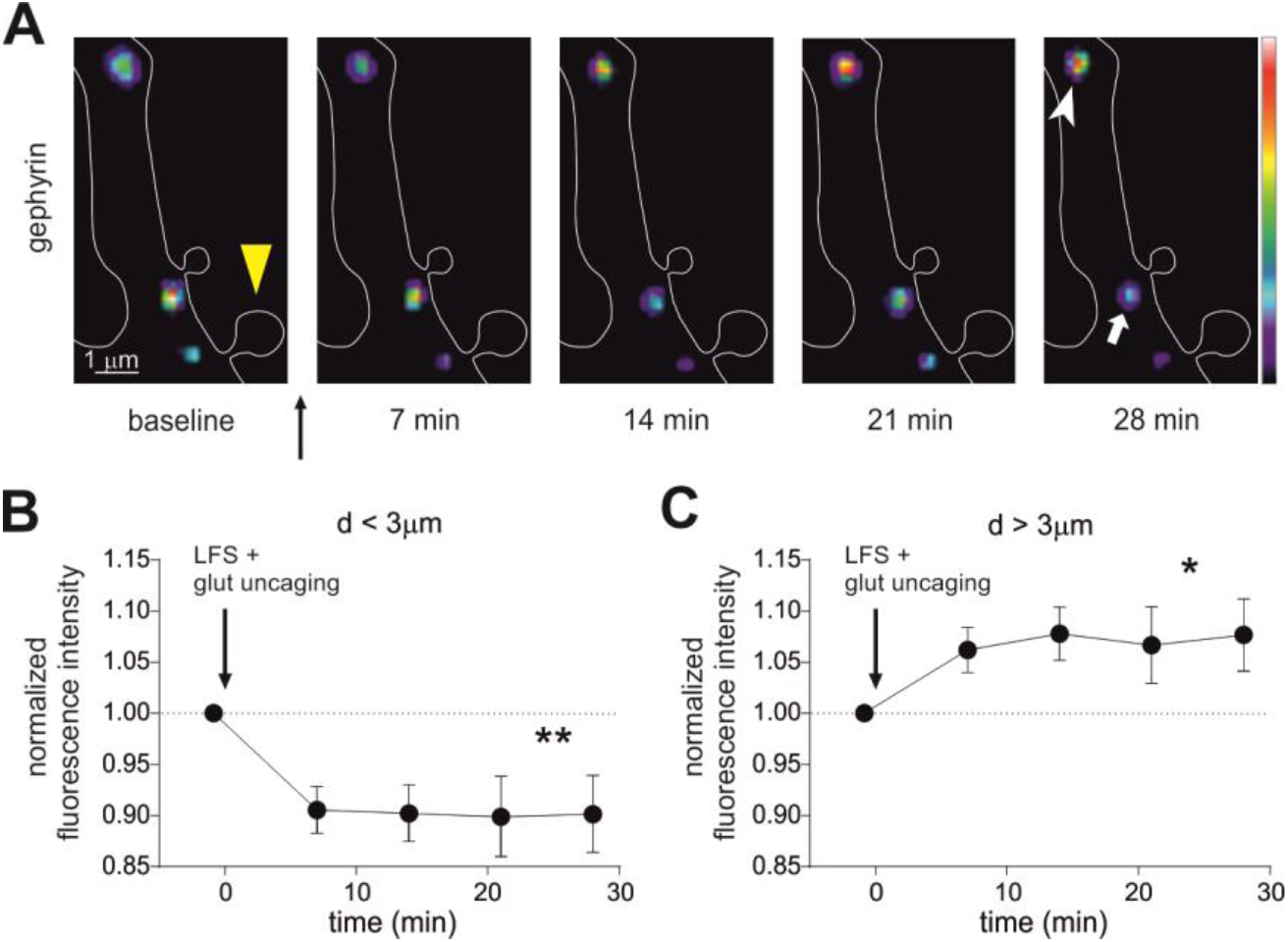
Gephyrin dynamics after single-spine LTP protocol. **A.** Representative dendritic portion of a Homer1c-DsRed expressing neuron (white outline) showing pseudocolored FingR-gephyrin fluorescent clusters at different time points before (baseline) and after the delivery of the LFS+glutamate uncaging protocol (yellow arrowhead). Clusters at distance > 3 μm (white arrowhead) from the stimulated spine (yellow arrowhead) were potentiated, whereas clusters at distance < 3 μm (white arrow) from the stimulated spine were depressed. Scale bar, 1 μm. **B.** Summary of the relative changes (after/before) in gephyrin fluorescence intensity quantified in clusters located at d < 3μm from the stimulated spine (n = 13, F_4,48_ = 4.8, p=0.02, RM one-way ANOVA followed by Dunnett’s post-test). **C.** Summary of the relative changes in gephyrin fluorescence intensity quantified in clusters located at d > 3μm from the stimulated spine (n = 13; F_4,48_ = 2.7, p = 0.04, RM one-way ANOVA followed by Dunnett’s post-test). Data are presented as mean ± SEM. *p < 0.05, **p < 0.01.

### Surface dynamics of GABA receptors after induction of single spine LTP

Postsynaptic long-term potentiation and long-term depression are associated with decreased or increased lateral mobility of postsynaptic receptors, respectively (Choquet and Triller, 2013; Petrini and Barberis, 2014). We next aimed to understand whether the opposing gephyrin modifications at increasing distances from a single spine expressing LTP is accompanied by differential surface dynamics of GABAA receptors (GABAARs). For this purpose, the lateral mobility of α1 subunit-containing GABAARs was analyzed by quantum dots-based single particle tracking (SPT) (see STAR Methods). In particular, with paired observations before and after the expression of single-spine LTP, we monitored the mobility of synaptic receptors at GABAergic synapses located either in the dendrite at a distance of ± 3 μm from an individual potentiated glutamatergic spine or further away (Figure 7A). Interestingly, at inhibitory synapses located > 3 μm from the potentiated spine – that is, those exhibiting iLTP (Figure 4B and S2B) – GABAARs were less mobile after the stimulating protocol, indicated by a reduced paired diffusion coefficient (before = 0.013 μm^2^s^-1^ and interquartile range (IQR) 0.008 - 0.029; after = 0.006 μm^2^s^-1^ and IQR: 0.002 - 0.011; n = 31 trajectories from 9 neurons, p<0.001; Figure 7B), an increased immobile fraction (before = 0.29 ± 0.07; after = 0.58 ± 0.07; n = 31, p<0.001; Figure 7C) and prolonged time spent at synapses (before = 36% ± 5%; after = 61% ± 6%; n=31, p=0.002; Figure 7D). We next considered GABAARs diffusing at synapses within a 3 μm range from the potentiated spine (d < 3 μm). Before the protocol, they exhibited diffusive properties comparable to more distant GABAARs (n_d>3_ = 31; n_d<3_ = 9, p>0.05; Figure 7B-7D). After the stimulation, those GABAARs close to the potentiated spine (d < 3 μm), (i.e., exhibiting iLTD in response to the single-spine LTP protocol, see Figure 4B and S2B), displayed markedly increased mobility (Figure 7A) as quantified in the diffusion coefficient (before = 0.012 μm^2^s^-1^ and IQR: 0.007 - 0.017; after = 0.022 μm^2^s^-1^ and IQR: 0.017 - 0.030; n = 9 trajectories from 5 neurons, p=0.04; Figure 7B) and immobile fraction (before = 0.29 ± 0.12; after = 0.04 ± 0.03; n=9, p=0.04; Figure 7C) after the stimulation. As expected, in these conditions of increased mobility, GABAARs escaped more frequently from the synaptic area, thus depleting inhibitory synapses of GABAARs during local iLTD (−61 % ± 11, n = 19; p<0.001; Figure 7E). As a control, we quantified the fraction of residual GABAARs at synapses within 3 μm of non-photo-stimulated spines at the end of each experiment (−6 % ± 3, n = 12; p = 0.25; Figure 7E). The negligible variation in synaptic GABAAR number in this control suggests that GABAARs selectively disperse from inhibitory synapses during local iLTD. However, the few GABAARs that remained at synapses during iLTD spent the same time in that compartment before and after single spine LTP induction (before = 39% ± 6%; after = 34% ± 8%; n = 9, p=0.49; Figure 7D). In line with these data, following the single spine LTP protocol, the steady state of the mean square displacement vs time curve (MSD) for GABAARs at synapses far from the potentiated spine (d > 3 μm) was reduced, thus indicating higher receptor confinement (n = 19 from 8 neurons, p=0.01; Figure 7F, left). In contrast, after the delivery of the same plasticity induction protocol, GABAARs in the dendritic range of ± 3 µm from the potentiated spine were less confined, as indicated by an increased MSD steady state (n = 9 from 5 neurons, p=0.01; Figure 7F, right).

**Figure 7:**
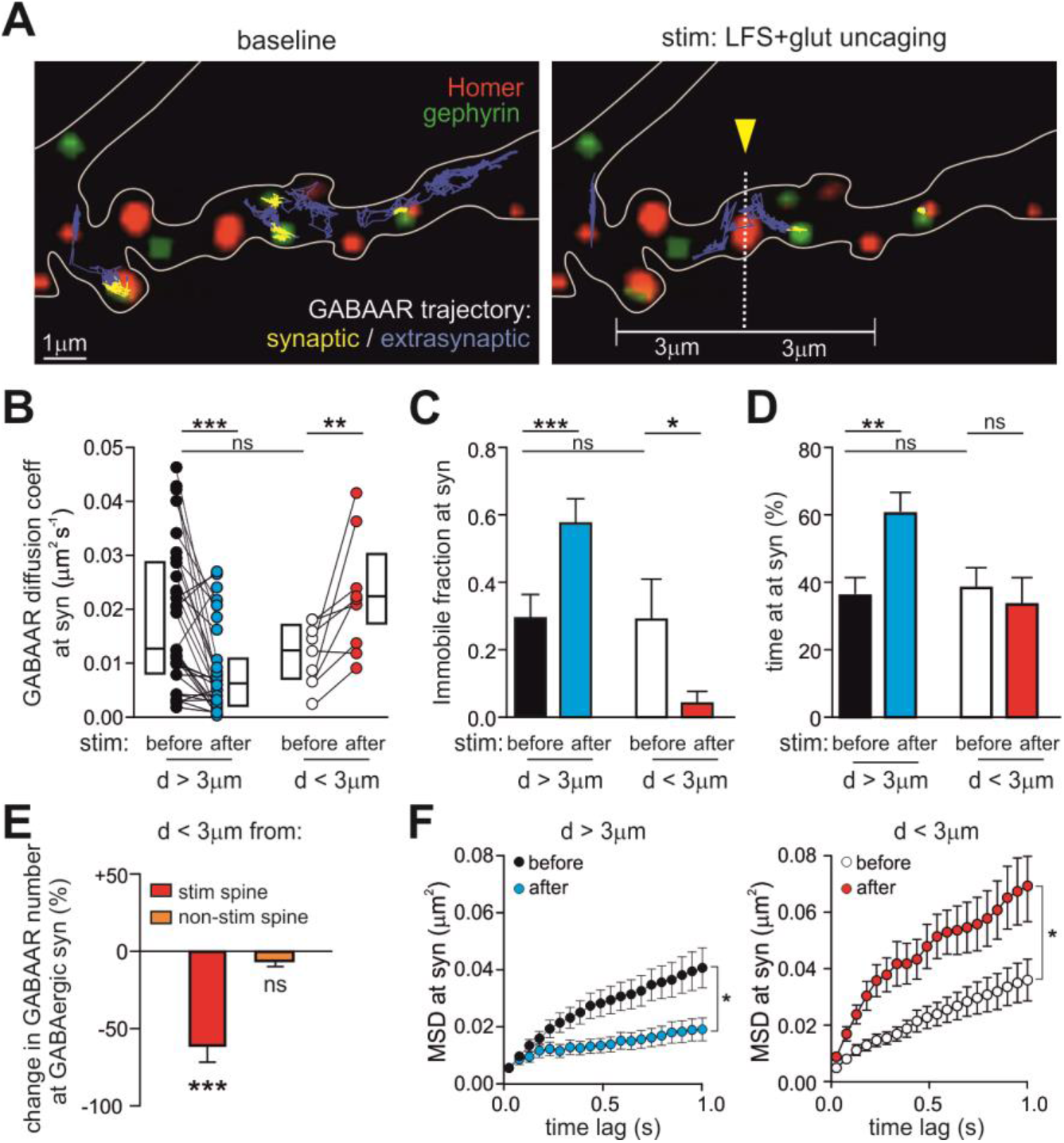
GABAA receptor lateral diffusion after the single-spine LTP protocol. **A.** Representative synaptic (yellow) and extrasynaptic (blue) trajectories of individual GABAARs diffusing on a gephyrin-GFP and Homer1c-DsRed expressing neuron, before (left) and after (right) the delivery of the LFS + MNI-glutamate uncaging protocol at the indicated spine (yellow arrowhead). Scale bar, 1 μm. **B-D.** Effect of LFS+glut uncaging on the surface mobility of GABAA receptors at synapses located at d > 3 or d < 3 µm from the potentiated spine. **B.** Paired diffusion coefficient values of synaptic GABAARs before and after the stimulating protocol (d > 3 µm: n = 31 trajectories from 9 neurons, p < 0.001, paired Wilcoxon test; d < 3 µm: n = 9 trajectories from 5 neurons, p = 0.04, paired Wilcoxon test). Comparison ‘before stim” d > 3 µm vs d < 3 µm: p = 0.19, Mann-Whitney test. **C.** Immobile fraction of synaptic GABAARs before and after the stimulating protocol (d > 3 µm: n = 31 from 9 neurons, p < 0.001, paired Wilcoxon test; d < 3 µm: n = 9 from 5 neurons, p = 0.03, paired Wilcoxon test). Comparison before d > 3 µm vs d < 3 µm: p = 0.62, Mann-Whitney test. **D.** Percentage of time spent at synapses of synaptic GABAARs before and after the stimulating protocol (d > 3 µm: n = 31, p = 0.003, paired Wilcoxon test; d < 3 µm: n = 9, p = 0.50, paired Wilcoxon test. Comparison ‘before stim” d > 3 µm vs d < 3 µm: p = 0.54, Mann-Whitney test. **E.** Variation in the number of synaptic GABAARs at GABAergic synapses close (d < 3 µm) to the potentiated spine (red; n = 19, p < 0.001, paired Wilcoxon test) or to a spine receiving the same protocol in the absence of MNI-glutamate (orange; n = 12, p = 0.25, paired Wilcoxon test). **F.** MSD versus time values of matched observations of individual GABAARs localized at d > 3μm (left) and d < 3μm (right) from the potentiated spine, before and after LFS+glut uncaging (d > 3 µm: n = 19 from 8 neurons, F_1,36_ = 7.0, p = 0.01; d < 3 µm: n = 9 from 5 neurons, F_1,16_ = 8.5, p = 0.01; RM two-way ANOVA followed by Bonferroni’s multiple comparison test). Unless otherwise stated, values are expressed as mean ± SEM.*p < 0.05, **p < 0.01, ***p < 0.001, ns = not significant. See also Figure S3.

It is worth noting that the single spine LTP protocol did not change the lateral diffusion properties of extrasynaptic receptors at distances > 3 μm (Figure S3A and S3B). Likewise, matched observations of individual extra-synaptic GABAARs in the range of 3 μm (d < 3 μm) from the potentiated spine showed unchanged diffusion coefficients and immobile fractions before and after the single spine LTP protocol (Figure S3C and S3D), while the percentage of time spent at the extrasynaptic domain increased (Figure S3C right). In order to rule out that the effect of the single spine LTP protocol on GABAAR diffusion was due to UV laser illumination, but instead required MNI-glutamate uncaging, we performed a control experiment in which the same protocol was performed without puffing MNI-glutamate (i.e., LFS paired with UV illumination). Synaptic GABAARs close to the illuminated spine (d < 3 μm) showed reduced mobility (before = 0.017 μm^2^s^-1^; IQR: 0.006 – 0.025; after = 0.005 μm^2^s^-1^ ; IQR: 0.004 - 0.09; n = 7 from 4 neurons; p=0.01; Figure S3E), increased immobile fraction (before = 0.22 ± 0.13; after = 0.62 ± 0.16; n=7; p=0.03; Figure S3E) and enhanced confinement (n = 4, p<0.001; Figure S3F), similarly to GABAARs at d > 3 μm from the potentiated spine, i.e. during iLTP (compare with Figure 7B-7D). Therefore, UV illumination (without MNI-glutamate) is not sufficient to reproduce the modifications of GABAAR dynamics observed during local iLTD. Overall, these results show that following the induction of single spine LTP, the spatial dependence of dendritic lateral diffusion of GABAAR faithfully corresponds with modulation of IPSC amplitude and gephyrin synaptic clustering.

## Discussion

In the present study, we investigated the interplay between excitatory and inhibitory synaptic plasticity in dendrites of hippocampal neurons. This is the first study analyzing how the expression of glutamatergic plasticity induced at individual glutamatergic spines affects the long-term plasticity of neighboring GABAergic synapses. Our findings indicate the existence of dendritic plasticity microdomains where the relative strength of excitatory and inhibitory synapses is oppositely regulated. In particular, we show that the application of non-Hebbian stimulations (postsynaptic delivery of LFS) concomitantly induces the depression of excitation (LTD) and the potentiation of inhibition (iLTP). Similarly, in conditions of Hebbian-like stimulation (LFS paired with MNI-glutamate uncaging), the potentiation of a single glutamatergic synapse (spine LTP) can be maximized by the surrounding depression of inhibition within a range of approximately 3 μm (local iLTD), whereas at dendritic sites far from the stimulated spines, non-Hebbian dendritic LTD and iLTP occur.

Our experiments indicate that calcium is fundamental for the induction of both the dendritic iLTP and local iLTD expressed near a potentiated spine. Indeed, LFS alone, which induces dendritic iLTP, causes a homogeneous dendritic calcium increase, which is likely sustained by back-propagating action potentials. The pairing of LFS with repetitive MNI glutamate uncaging, responsible for the local iLTD, induced a further dendritic calcium increase that was confined within a range of 3-4 μm from the stimulated spine. This local additional calcium increase could arise from the activation of dendritic voltage gated calcium channels (VGCCs), although it is not clear whether the depolarization reached in the shaft near the spine is sufficient to activate VGCCs, given the heavy filtering of synaptic potentials by the spine neck (Yuste, 2013). In addition, calcium would permeate through the spine neck, although both the buffering and the rapid extrusion of calcium would be expected to largely compartmentalize calcium at spines (Sabatini et al., 2002). Nevertheless, both VGCC activation and calcium permeation from the spine to the shaft might account for the dendritic local calcium increase during the repeated spine activation used to induce single spine LTP.

Previous work using chemical protocols to induce inhibitory plasticity show that sustained calcium influx leads to depression (Bannai et al., 2009, 2015; Muir et al., 2010), while mild calcium entry determines potentiation (Marsden et al., 2007; Marsden et al., 2010; Petrini et al., 2014, Chiu et al., 2018) of GABAergic synapses. This indicates that GABAergic plasticity might follow an opposite calcium rule with respect to that of glutamatergic synapses, where high and low intracellular calcium rises have been linked to expression of LTP and LTD, respectively (Coultrap et al., 2014; Lisman, 2001). Thus, a possible explanation for the paradigm of GABAergic plasticity expression shown here is that LFS alone (responsible for a mild calcium entry) could induce iLTP, while the summation of calcium increase due to the pairing of LFS with single spine MNI-glutamate uncaging would locally induce high calcium concentrations sufficient to reach the threshold for iLTD.

We propose that that such a high local calcium concentration in the shaft near the potentiated spine activates locally available calpain, which in turn promotes local iLTD through the proteolysis of gephyrin and the consequent destabilization of the inhibitory postsynaptic density (iPSD), a mechanism already demonstrated by independent studies (Costa et al., 2016; Tyagarajan et al., 2013). Since calpain is widely distributed both in spines and dendritic shafts, it is also possible that following the pairing of LFS with focal glutamate uncaging, activated spine calpain would diffuse to the parent dendrite, similarly to several other signaling molecules responsible for short-range heterosynaptic interaction among glutamatergic synapses (Chen and Sabatini, 2012; El-Boustani et al., 2018; Harvey et al., 2008; Murakoshi et al., 2011; Oh et al., 2015; Yasuda, 2017).

We found that following the induction of single-spine LTP, the spatially-regulated, opposite uIPSC plasticity at different distances from the potentiated spine was paralleled by changes of gephyrin aggregation. Indeed, imaging experiments showed that gephyrin clustering was reduced within 3 μm of the potentiated spine (where local iLTD was expressed), while it was increased at d > 3 μm from the potentiated spine (where the iLTP was observed). These data are in line with the notion that gephyrin accumulation at the iPSD contributes to clustering GABAARs, thus representing a proxy for the amplitude of GABAergic synaptic currents (Bannai et al., 2009; Battaglia et al., 2018; Flores et al., 2015; Petrini et al., 2014; Tyagarajan et al., 2011; Villa et al., 2016). However, it must be stressed that given the rich diversity of GABAergic synaptic proteins, both the involvement of gephyrin and the mechanisms of plasticity observed here could significantly vary at synapses located in specific neuron types and neuronal subregions along the axo-dendritic axis (Chiu et al., 2019; Fritschy et al., 2012). In addition to gephyrin clustering, the opposite regulation of GABAergic plasticity also entailed an opposite modulation of surface GABAAR lateral diffusion. Indeed, following the induction of single spine LTP, we observed GABAAR immobilization at potentiated inhibitory synapses far from the potentiated spine (d > 3 μm, iLTP), whereas in the vicinity of the stimulated spine (d < 3 μm, local iLTD), GABAAR mobility increased, as did consequent receptor dispersal from inhibitory synapses. This is consistent with previous work demonstrating that sustained calcium entry increases GABAAR mobility, while moderate calcium entry reduces it (Bannai et al., 2009, Petrini et al., 2014; Bannai et al., 2015). Overall, we find coherent spatial regulation of gephyrin clustering and GABAA receptor mobility that sustains the expression of iLTP or local LTD as a function of local unevenness in dendritic calcium changes upon single-spine LTP. However, although it has been shown that calcium-dependent signaling modulates the interaction among gephyrin, GABAA receptors and other synaptic inhibitory scaffold proteins to regulate iPSD clustering (Barberis, 2019; Chiu et al., 2019; Panzanelli et al., 2011; Tyagarajan and Fritschy, 2014), the precise interplay of these molecular players during synaptic plasticity remains to be fully elucidated.

The main finding of our study – that excitatory plasticity at an individual spine can affect neighboring GABAergic synapses in the range of few microns – is in good agreement with previous studies that tackled the issue of interplay between dendritic synapses. The in vivo dynamic changes of excitatory and inhibitory synapses in pyramidal neurons of the visual cortex over days after monocular deprivation (MD) were clustered in dendritic sub-regions of ∼ 10 μm (Chen et al., 2012), indicating that glutamatergic and GABAergic synapses during plasticity are locally coordinated. Similarly, it has been recently demonstrated in vivo in dendrites of visual cortex (V1) neurons that the Hebbian potentiation of a specific spine through pairing of visual stimuli and optogenetic activation leads to selective depression of neighboring spines (El-Boustani et al., 2018). Similarly, the potentiation of an individual spine or a pair of spines through either high frequency stimulation or motor skill training induces the shrinkage of adjacent spines. Along the same lines, the potentiation of an individual spine lowers the threshold for the potentiation of neighboring spines (Harvey and Svoboda, 2007). These studies indicate that the fundamental units of dendritic synaptic plasticity are clusters of synapses located on the same dendritic portion rather than individual synapses. This concept has been also explored by studies exploiting modelling approaches (Larkum and Nevian, 2008; Rabinowitch and Segev, 2008). Thus, our data converge to the hypothesis of clustered dendritic synaptic plasticity and represent a piece of evidence that plasticity microdomains are formed by both glutamatergic and GABAergic synapses. Recently, in vivo studies in visual cortex have shown that over the period of several days, GABAergic dendritic synapses can be frequently formed and eliminated in the same location. This has led to a new definition of inhibitory synaptic plasticity that relies on substantial dynamic rearrangements of GABAergic synapses (Chen et al., 2012; Villa et al., 2016). In the light of this new perspective, it remains to be established whether the local plasticity interplay documented here in its early phase (30 min) is preserved in the long-term or rather represents a transient phase of GABAergic synaptic dynamism, e.g., after days post-plasticity induction.

A previous electron microscopy study investigated the ultrastructural changes of both glutamatergic and putative inhibitory synapses before and after the induction of synaptic plasticity through the delivery of theta burst stimulations (TBS) (Bourne and Harris, 2011). TBS increases the size of both excitatory and inhibitory dendritic synapses at the expense of their density, keeping constant the total areas of synaptic contacts. Such TBS-induced parallel changes at glutamatergic and GABAergic synapses differ from our results (whereby excitatory and inhibitory synapses respond with opposite molecular changes to the same stimulus), although it is not clear whether the TBS-induced modifications of inhibitory synapses result in the alteration of inhibitory synaptic strength. In more recent work, TBS has been reported to potentiate IPSCs, through the increase of both the size and the density of inhibitory synapses (Flores et al., 2015). Thus, although both the LFS presented here and TBS are able to induce GABAergic long-term potentiation, opposite results have been reported for the concomitant changes of glutamatergic synaptic plasticity (Bourne and Harris, 2011). This exemplifies the importance of simultaneously considering both excitatory and inhibitory synapses when studying the net plasticity output in response to specific plasticity-induction paradigms. With this in mind, we show that in both non-Hebbian and Hebbian stimulation conditions, synaptic excitation and inhibition are always oppositely regulated. This result could appear to contrast with a previous study showing that glutamatergic and GABAergic inputs in principal neurons in layer 2/3 of visual cortex are remarkably balanced (Xue et al., 2014). However, in our model, local synaptic E/I imbalance could still be equalized in larger portions of dendrites or in the whole neuron, where equilibrium among the aforementioned opposite plasticity domains could be reached. This highlights the concept that the synaptic E/I balance, known to be altered in neurological disorders (Antoine et al., 2019; Lewis et al., 2012; Rubenstein and Merzenich, 2003) could be selectively disrupted in specific dendritic micro-domains, even where the overall E/I balance is preserved at the level of the whole neuron. Such plasticity interplay of glutamatergic and GABAergic synapses in microscale compartments is expected to significantly impact dendritic signal integration and processing, in particular in view of the compartmentalization of both excitatory and inhibitory synaptic inputs in dendritic sub-regions of pyramidal neurons(Klausberger and Somogyi, 2008; Spruston, 2008).

## ACKNOWLEDGMENTS

We thank Stefania Guazzi, Mattia Pesce, Alice Gino and the IIT animal facility for technical help. This work has been supported by Telethon-Italy (GGP11043) and Compagnia di San Paolo (ROL-4318).

## AUTHOR CONTRIBUTION

Conceptualization, A.B.; Methodology, T.R., M.R., E.M.P. and A.B.; Validation, T.R., M.R., E.M.P. and A.B.; Formal Analysis, T.R., M.R., E.M.P. and A.B.; Investigation, T.R. and M.R.; Resources, T.R., M.R. and A.P.; Data Curation, T.R., M.R.; Writing – original Draft, A.B., M.R. and T.R.; Writing – Review & Editing, A.B. and E.M.P.; Visualization, T.R., E.M.P., M.R., A.P., and A.B.; Supervision, A.B.; Project Administration, A.B.; Funding Acquisition, A.B.

## DECLARATION OF INTEREST

The authors declare no competing interests

## METHODS

### CONTACT FOR REAGENT AND RESOURCE SHARING

Further information and requests for resources and reagents should be directed to and will be fulfilled by the Lead Contact, Andrea Barberis.

### EXPERIMENTAL MODEL AND SUBJECT DETAILS

#### Animals

All the experiments were carried out in accordance with the laws of Italian Ministry of Health and the guidelines established by the European Communities Council (Directive 2010/63/EU, 2010). Parvalbumin-tdTomato (PV-tdTomato) mice were obtained at the IIT animal facility by breeding Ai9 mice with PVCRE mice. Ai9 (B6.Cg-*Gt(ROSA)26Sortm9(CAGtdTomato) Hze*/J – Jackson Laboratory, USA) mice carrying a *loxP*-flanked STOP cassette, that prevents the transcription of a CAG promoter-driven red fluorescent protein variant (tdTomato) were used as a Cre reporter strain. PVCRE (B6;129P2-Pvalbtm1(cre)Arbr/J - Jackson Laboratory, USA) mice express the Cre recombinase in Parvalbumin-expressing interneurons without disrupting the endogenous Parvalbumin locus (Pvalb) expression. The resulting offspring PV-td tomato has the STOP cassette removed in Parvalbumin-potitive interneurons and the consequent expression of tdTomato.

#### Primary neuronal cultures

Cultures of hippocampal neurons were prepared from P1-P3 Parvalbumin-tdTomato mice of either sex using a previously published protocol (de Luca et al., 2017) modified from (Baudouin et al., 2012). Briefly, hippocampi were dissected, quickly sliced and digested with trypsin in the presence of DNAase, mechanically triturated, centrifuged at 80g and re-suspended. Neurons were plated at a density of 90 x 10^3^ cells/ml on poly-D-lysine (0.1 μg/ml) pre-coated coverslips. Cultures were kept in serum-free Neurobasal-A medium (Invitrogen, Italy) supplemented with Glutamax (Invitrogen, Italy) 1%, B-27 (Invitrogen, Italy) 2% and Gentamycin 5 µg/ml at 37°C in 5% CO2 up to 30 days in vitro (DIV). During this period, half of the medium was changed weekly. Experiments were conducted at DIV 15-27.

### Plasmid constructs

Enhanced GFP (eGFP) was expressed from the pEGFP-N1 (Clontech). Homer1c-DsRed and Homer1c-GFP plasmids encoding for Homer1c fused with DsRed and GFP at the N terminus, respectively were kindly provided by Dr. D. Choquet (Petrini et al., 2009). EGFP-gephyrin was a gift from Prof. E. Cherubini. FingR-gephyrin-GFP (received from Dr C. Duarte) was expressed from pCAG_GPHN.FingR-eGFP-CCR5TC, a plasmid encoding for FingRs (Fibronectin intrabodies generated with mRNA display), that bind endogenous gephyrin with high affinity and allow the visualization of gephyrin clusters using GFP as a reporter (Gross et al., 2013). GABAA receptor α1 subunit carrying the Hemagglutinin (HA) tag between the IV and V amino acid of the mature protein has been described previously (de Luca et al., 2017)

### METHOD DETAILS

#### Transfection and synapse visualization

Neurons were transfected with either using Effectene (*Qiagen*, Germany) at 6-7 days in vitro (DIV) or Lipofectamine 2000 (Thermofisher) 24/72 hours before the experiments, following the companies’ protocols. All experiments were performed from 14 DIV to 21 DIV. In most experiments, excitatory and inhibitory synapses were visualized by transfecting Homer1c-DsRed and gephyrin-EGFP, respectively. GABAergic synapses were also identified by live immunolabelling of the presynaptic marker vGAT using the anti-vGAT-Oyster550 antibody (Synaptic Systems, Germany) which is directed against the luminal part of the protein, diluted in culture medium and incubated for 30 min at 37°C.

#### Antibodies and drugs

Anti-vGAT-Oyster 550 antibody was purchased from Synaptic System (Goettingen, Germany). Anti-HA antibody was from Roche (Milan, Italy). BAPTA (1,2-bis(o-aminophenoxy)ethane-N,N,N’,N’-tetraacetic acid), L-NAME (L-NG-Nitroarginine methyl ester), Nifedipine (1,4-Dihydro-2,6-dimethyl-4-(2-nitrophenyl)-3,5-pyridinedicarboxylic acid dimethylester), and Bicuculline were purchased from Sigma (Milan, Italy). KN-93 and KN-92 were acquired from Millipore Merck (Darmstadt, Germany). APV (D-(-)-2-Amino-5-phosphonopentanoic acid), CNQX (6-Cyano-7-nitroquinoxaline-2,3-dione), ω-conotoxin MVIIC, ω-conotoxin GVIA, DPNI-caged-GABA (1-(4-Aminobutanoyl)-4-[1,3-bis(dihydroxyphosphoryloxy)propan-2-yloxy]-7-nitroindoline) and MNI-caged-L-glutamate ((S)-α-amino-2,3-dihydro-4-methoxy-7-nitro-δ-oxo-1H-indole-1-pentanoic acid) were purchased from Tocris (Bristol, UK). Rhod-2 tripotassium salt was purchased from AAT Bioquest (Sunnyvale, CA, USA).

#### Electrophysiological recordings

Inhibitory and excitatory postsynaptic currents (IPSCs and EPSCs, respectively) were recorded at room temperature in the whole-cell configuration of the patch-clamp technique. External recording solution contained (in mM): 145 NaCl, 2 KCl, 2 CaCl_2_, 2 MgCl_2_, 10 glucose, and 10 HEPES, pH 7.4. Patch pipettes, pulled from borosilicate glass capillaries (Warner Instruments, LLC, Hamden, USA) had a 4 to 5 MΩ resistance when filled with intracellular solution. In all experiments with the exception of paired-patch electrophysiological recordings, the intracellular solution contained (in mM): 10 KGluconate, 125 KCl, 1 EGTA, 10 HEPES, 5 Sucrose, 4 MgATP (300mOsm and pH 7.2 with KOH). Paired-patch recordings were performed with an intracellular solution containing (in mM): 130 KGluconate, 20 KCl, 1 EGTA, 10 HEPES, 5 Sucrose, 4 MgATP (300mOsm and pH 7.2 with KOH). In a subset of paired-patch recordings 20 KCl was replaced with 5 KCl. Since the use of these two intracellular solutions gave comparable results, data were merged. In the paired-patch experiments using BAPTA, 1mM EGTA was replaced with 11mM BAPTA in the presence of 120 mM KGluconate. Currents were recorded using Clampex 10.0 software (Molecular Devices, Sunnyvale, CA). The stability of the patch was checked by monitoring the input resistance during the experiments to exclude cells exhibiting more than 15% changes from the analysis. Currents were sampled at 20 KHz and digitally filtered at 3 KHz using the 700B Axopatch amplifier (Molecular Devices, Sunnyvale, CA). IPSCs and EPSCs were analyzed with Clampfit 10.0 (Molecular Devices, Sunnyvale, CA).

In paired-patch experiments, a small current was injected into presynaptic neurons in the current clamp mode to keep their membrane potential close to -65 mV. Action potentials, evoked in the presynaptic neuron by injecting depolarizing current pulses (0.8-1 nA for 5-7 ms) at a frequency of 0.1 Hz, elicited IPSCs or EPSCs that were recorded from the postsynaptic neuron voltage-clamped at -65 mV. When the paired-patch involved a presynaptic PV+ interneuron and a putative pyramidal neuron, GABAergic IPSCs were pharmacologically isolated by the continuous perfusion of CNQX (10 µM) to prevent glutamatergic synaptic transmission. When the presynaptic and postsynaptic neurons were two putative pyramidal neurons, EPSCs were isolated by the continuous perfusion of Bicuculline (10 µM) to prevent GABAergic transmission. IPSCs or EPSCs were continuously acquired from 5 min before to 30 min after the delivery of the electrical plasticity-inducing protocol (see below). IPSCs and EPSCs data were binned in 1 min intervals and normalized to the mean of the baseline amplitude. Data are expressed as normalized values after/before. In the text, we report stimulation-induced average changes in current amplitude between 25 and 30 min after the protocol and expressed as fold-change of the baseline.

#### Neurotransmitter Uncaging

GABA and Glutamate were photoreleased from DPNI-GABA and MNI-glutamate after illumination by a 378 nm diode laser (Cube 378, 16 mW, Coherent Italia, Italy). MNI-glutamate (5 mM) or DPNI-GABA (1 mM) were dissolved in extracellular solution and locally applied near the synapses through a pulled glass capillary (2-4 µm tip diameter) placed at 10-30 µm in the x-axis and at 5-10 µm in the z-axis from the region of interest (ROI), using a pressure-based perfusion system (5/10 psi) (Picospritzer, Parker, USA). The laser beam was focused on the sample by means of an Olympus Apo-plan 100X oil-immersion objective (1.4 NA). A beam expander was placed in the optical path between the laser source and the objective in order to achieve a complete filling of the objective pupil, a conditions that maximizes the focusing capability of the objective, thus minimizing the spot size on the sample. The measured point spread function (PSF) had a lateral dimension of 487±55 nm (FWHM, n = 6). The laser beam was steered in the field of view by using a galvanometric mirrors-based pointing system able to illuminate specific regions of interest outlined around glutamatergic and GABAergic synapses defined by Homer1c-DsRed and gephyrin-GFP (UGA32, Rapp OptoElectronics, Hamburg, Germany). Synchronization of optical uncaging and electrophysiological recordings was controlled with the UGA32 software interfaced with the Clampex 10.0 software (Molecular Devices, Sunnyvale, CA, USA). Both MNI-glutamate and DPNI-GABA uncaging currents (uEPSCs and uIPSCs, respectively) were elicited by 500-1000 µs laser pulses with a power intensity of 80-100 µW at the exit of the objective. In double-uncaging experiments, the same uncaging settings were applied, with MNI-glutamate and DPNI-GABA loaded in two glass capillaries independently positioned in the ROIs and independently controlled by the aforementioned pressure-based perfusion system. The time course of uncaging current amplitude changes upon plasticity induction was quantified by binning data in 10 min intervals and by normalizing them to the mean of the amplitude at baseline time points. In the text, we report the values of stimulation-induced average changes in current amplitude at 27 min after the protocol expressed as fold-change of the baseline.

#### Plasticity induction

The non-Hebbian plasticity-inducing protocol consisted of action potential (AP) trains elicited in the postsynaptic neuron at 2 Hz for 40 seconds (low frequency stimulation, LFS) in the current clamp configuration. AP was elicited by the injection of depolarizing current pulses (0.8-1 nA for 5-7 ms) (0.8-1 nA for 5-7 ms). Single spine LTP (for a Hebbian stimulation) was induced by pairing the aforementioned LFS with repetitive MNI-glutamate uncaging at 4 Hz at individual spines for 40 seconds (see Neurotransmitter Uncaging). In the text, this protocol has been referred to also as “LFS + MNI-glutamate uncaging”. Experiments aimed at identifying the contribution of i) different calcium sources ii) CaMKII role or iii) nitric oxide (NO) role in inhibitory plasticity were performed in the same conditions described above during the bath application of APV (50 µM), L-NAME (50 µM), KN-93 (5 µM), KN-92 (5 µM), ω-conotoxin MVIIC (2 µM), ω-conotoxin GVIA (3 µM) or Nifedipine (10 µM) as described in the text. In the experiments performed to study the involvement of the calpain, neurons were preincubated with the calpain inhibitor III MDL28170 (50 µM) for 30 minutes before the delivery of the single spine LTP protocol.

#### Live-Cell Imaging

Hippocampal primary cultures from PV-tdTomato mice were transfected with FingR-gephyrin-GFP or gephyrin-GFP. Samples were illuminated with a LED light source (Spectra X, Lumencor) through 475/34 nm and 543/22 filters (Semrock, Italy). GFP and tdTomato fluorescence was detected using a 520/35 nm and 593/40 nm filters respectively (Semrock, Italy). Neurons positive for GFP were identified, patched and stimulated with the both the non-Hebbian and Hebbian electrophysiological plasticity-inducing protocol. Neurons positive for both GFP and PV-tdTomato were excluded. Images were acquired with the digital camera Hamamatsu, EM-CCD C9100 mounted on a wide field inverted fluorescence microscope (Nikon Eclipse Ti) equipped with an oil-immersion 60X (1.4 NA) or with the digital camera EM-CCD Photometric QuantEM:512SC mounted on a wide field inverted fluorescence microscope (Olympus IX 70) equipped with an oil immersion 100X objective (1.4 NA), for the imaging of gephyrin-GFP or FingR-gephyrin-GFP clusters, respectively. Acquisition and quantification of gephyrin clusters fluorescence were performed by using the MetaMorph 7.8 software (Molecular Devices).

Images of FingR-gephyrin-GFP or gephyrin-GFP clusters fluorescence was acquired before and after (up to 30 minutes) the application of the LFS protocol (non-Hebbian stimulation) or the LFS+MNI-glutamate uncaging protocol at individual spines (Hebbian stimulation). Focal plane was set by the operator and maintained fixed for the duration of the experiment. Gephyrin clusters that changed their focal plane after the delivery of the stimulation, were discarded from the analysis. The same light exposure time was used for the acquisition of all images and was set to avoid signal saturation. After background correction, a user-defined intensity threshold was applied to the maximal projection of each image-stack to create a binary mask for the identification of gephyrin clusters. For the analysis of gephyrin clusters, regions were created around each cluster in the binary mask after 2 pixel enlargement. As such, we aimed at avoiding the possibility of underestimating gephyrin fluorescence over time due to the changes in the cluster size/position after the delivery of the protocol. Average fluorescence intensity of each cluster was measured and normalized to control experiments in which the stimulation was omitted.

#### Calcium imaging

Calcium imaging experiments were performed by using Rhod-2 (Minta et al., 1989). The rationale for the choice of this red shifted rhodamine-based calcium indicator with respect to the more commonly used green-emitting indicators was to maximize the separation between the wavelength of the laser used for neurotransmitter uncaging (378 nm) and the indicator absorption spectrum, thus minimizing the possible photobleach of the indicator. Previous studies have shown that the positive net charge of the Rhod-2 molecule favors intracellular Rhod-2 accumulation in mitochondria (Collins et al., 2001). However, this particular Rhod-2 partitioning between cytosol and mitochondria has been mainly observed with the cell permeant form of Rhod-2 (Rhod-2 AM). In contrast, the cell-impermeant form has been used to record bona fide cytosolic calcium in electrophysiological studies (Kaiser et al., 2004; Yasuda et al., 1998). In our calcium experiments with Rhod-2, we observed that, while it efficiently dialyzed in dendrites, it showed limited diffusion into spines. However, since our goal was to study calcium dynamics in the dendritic shaft, we reasoned that such Rhod-2 feature could contribute to maintain unperturbed the spine calcium dynamics, while recording the dendritic one.

Neurons were loaded with Rhod-2 (80 µM) through the patch pipette for at least 5 minutes after reaching the whole-cell configuration to allow the diffusion of Rhod-2 in proximal dendrites. Rhod-2 fluorescence was acquired with the digital EM-CCD QuantEM:512SC camera (Photometrics) mounted on a wide field inverted fluorescence microscope (Olympus IX 70) equipped with an oil-immersion 100X objective (1.4 NA) and the MetaMorph 7.8 software (Molecular Devices). The LFS paired with MNI-glutamate uncaging was delivered at individual spines (Hebbian stimulation). During this protocol, we recorded calcium dynamics in a dendritic region centered below the photostimulated spine. Concomitantly, calcium dynamics was also recorded in another region on a different dendritic branch of the same neuron (at a similar distance from the soma) centered below a reference, non-photostimulated spine. Since the latter region was distant from the potentiated spine, it was receiving only the LFS, so hereafter it will be referred to as “LFS” conditions. The onset of calcium responses recorded in the two regions reached plateau in a few seconds after stimulation. Thus, the stimulation protocol duration was reduced to 10 seconds (instead of the full-length stimulation of 40 seconds) in order to minimize fluorescence photobleaching. Therefore, the total duration of the recording was 16 seconds (i.e., 160 frames acquired at 10 Hz) including 3 seconds before (baseline), 10 seconds during and 3 seconds after (recovery) the stimulation protocol.

For the data analysis, we considered dendritic portions of 14 µm centered below the stimulated or the reference spine - which was usually chosen at approximately 10-30 µm from the soma. Every dendritic portion was sub-divided in 7 regions of interest (ROIs) of 2 µm length, with the central one being centered below the spine. The width of each region was adjusted to the thickness of the dendrite. In each region, changes in the Rhod-2 fluorescence intensities induced by the LFS or LFS + MNI-glutamate uncaging were calculated as ΔF/F_0_, where ΔF is the difference between the average fluorescence intensities at plateau and that before the delivery of the protocol. F_0_ is the average fluorescence intensity measured before the stimulation. In order to quantify calcium variations induced by the pairing of MNI-glutamate uncaging with respect to LFS alone, the ΔF/F_0_ recorded upon LFS+MNI-glutamate uncaging was normalized to that observed upon LFS (i.e., Figure 5D and 5E). When considering the spatial spread of calcium variations induced by the stimulating protocols (Figure 5E), the aforementioned normalization was computed for each ROI.

#### Quantum dot labelling and imaging

In the experiments aimed at monitoring the modulation of GABAA receptor lateral mobility during spatially-regulated synaptic plasticity, we combined SPT experiments with electrophysiology and plasticity induction (see sections above). Non-Hebbian or Hebbian stimulation protocols were delivered to neurons expressing the HA-tagged α1 subunit of GABAA receptor along with Homer1c-DsRed and gephyrin-GFP. The surface labelling of the HA tag with QDs allowed to selectively probe the mobility of GABAARs belonging to the neuron that received the plasticity protocol.

Before the experiment, QDs 655 (Invitrogen) were diluted in PBS and pre-exposed to casein 1X (Vectorlab, Italy) for 15 min to prevent QD non-specific binding. Then, living neurons were incubated with the anti-HA antibody (Roche) 1 μg/ml in the electrophysiology external recording solution for 4 minutes and subsequently with the diluted QDs solution for 3 minutes. The final concentration of QD was 0.1 nM. Control experiments omitting the anti-HA antibody were performed to validate the antibody-specific labelling of HA-tagged GABAARs.

SPT experiments were acquired by live-cell imaging on a wide field inverted fluorescence microscope (Olympus IX 70) equipped with a diode-based illumination device (Lumencor, SpectraX Light Engine, Optoprim, Italy), an EM-CCD camera (QuantEM:512SC, Photometrics, pixel size 16 μm) and an Apo-plan oil-immersion 100X objective 1.4 NA (Olympus). For each neuron, we chose a dendritic portion where we first localized glutamatergic and GABAergic synapses by Homer1c-DsRed and gephyrin-GFP fluorescence acquired with appropriate excitation and emission filter sets (ex: 543/22, 472/30, em: 593/40, 520/35, respectively) to achieve a 2D map of the relative localization of excitatory and inhibitory synapses. QD fluorescence acquired with specific filters (ex: 435/40 and em: 655/15 filters, Semrock, Italy) was monitored over time by recording movies of 600 consecutive frames at 20 Hz using the Metamorph 7.8 software (Molecular Devices, USA). The mobility of GABAAR-QD complexes was probed in the same field of view before and 30 minutes after the induction of synaptic plasticity, either with the LFS or with LFS paired with MNI-glutamate uncaging. During the experiments, neurons were kept at 28°C (TC-324B Warner Instrument Corporation, CT, USA) in an open chamber and continuously superfused with the recording solution at 12 ml/h.

#### Single particle tracking

Tracking of QD-labelled GABAAR was performed as previously described (Petrini et al., 2009; de Luca et at., 2017). The spatial coordinates of single QDs were identified in each frame as sets of > 4 connected pixels using two-dimensional object wavelet-based localization at sub-diffraction limited resolution (∼ 40 nm) using the MIA software which is based on simulated annealing algorithm. Continuous tracking between blinks was performed with an implemented version of custom software originally written in MATLAB (The Mathworks Inc., Italy) in Dr Choquet’s lab, based on a QD maximal allowable displacement (4 pixels) during a maximal allowable duration of the dark period. This stringent reconnection of trajectories across QD blinking combined with the highly diluted QD labelling have been set to avoid erroneous reconnection of neighbouring QDs in the same trajectory and to provide unambiguous observations of individual receptor-QD complex trajectories. When, occasionally, two QDs were too close to be unambiguously identified, they both were discarded from the analysis. Receptor trajectories were defined as synaptic (or extrasynaptic) when their spatial coordinates matched (or not) those of clustered gephyrin-GFP fluorescence. Although the definition of the synaptic compartments was diffraction limited, the sub-wavelength resolution of the single particle detection (∼40 nm) allowed accurate description of receptor mobility within such small regions. For each receptor-QD complex, the instantaneous diffusion coefficient, *D*, was calculated from the linear fits of the *n*=1–4 values of the MSD versus time plot, using a custom-made software developed by Dr Choquet (Bordeaux, France). For two-dimensional free diffusion, MSD is represented by the equation: MSD*(t)=<r^2^>=4Dt*.

*MSD(t)* was calculated according to the formula:

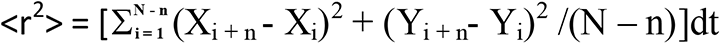

Only reconstructed trajectories with >80 frames were retained for the analysis. The diffusion coefficients are presented as median and IQR (i.e. the interquartile range) defined as the interval between 25–75% percentiles. The immobile fraction is defined as the relative duration of the residency of a receptor-QD complex in a given compartment with coefficient <0.0075 μm^2^ s^-1^. This threshold represents the local minimum of the bimodal distribution of synaptic GABA_A_R diffusion coefficients. To achieve a more complete characterization of GABAA receptor diffusion, we also measured the percentage of time spent by each receptor-QD in a given compartment (synaptic or extrasynaptic). In the case of local iLTD, when GABAAR disperse from inhibitory synapses, leaving few receptor-QD complexes for quantification, we also calculated the percentage of receptor number found at synapses after plasticity induction as compared to before the protocol.

### QUANTIFICATION AND STATISTICAL ANALYSIS

For each experiment quantifications and statistical details (statistical significance and test used) can be always found in the main text and figure legends. Unless otherwise stated, normally distributed data are presented as mean ± SEM (standard error of the mean), whereas non-normally distributed data are given as medians ± IQR (inter quartile range). For electrophysiological experiments in the paired-patch configuration as well as for gephyrin live-cell imaging and intracellular calcium imaging experiments n represents the number of neurons observed. In uncaging experiments, the number of synapses (n) is reported along with the number of neurons considered. For SPT experiments, n indicates the number of receptor trajectories, followed by the number of neurons observed. Each experiment was repeated on neurons obtained from at least three different cultures. The sample size used in each experiment was based on previous electrophysiological, live-cell imaging and SPT experiments (Petrini et al., 2014, de Luca et al., 2017). Data and statistical analysis was performed using Prism 5.0 and 6.0 Software (GraphPad Prism, USA). Normally distributed data sets were compared using the two-tailed unpaired Student’s t-test or, in the case of paired data, with the paired t-test. Non–Gaussian data sets were tested by two-tailed non-parametric Mann-Whitney U-test, or in the case of paired data, Wilcoxon paired test. In paired-patch experiments, statistical differences in time course data within a group was quantified by one-way ANOVA variance test followed by Turkey’s multiple comparison test. For time course of uncaging and imaging experiments exhibiting only one time point at the baseline, one-way ANOVAs were performed followed by Dunnett’s multiple comparisons test. When possible, RM ANOVA was used, as indicated. Statistical significance between more than two normally distributed data-sets was tested by two-way ANOVA variance test followed by Bonferroni’s multiple comparisons test. Indications of significance correspond to p-values as follows: p<0.05 (*), p<0.01 (**), p<0.001 (***) and non-significant (ns), i.e. p>0.05.

## Notes

### Competing Interest Statement

The authors have declared no competing interest.

