## Supplemental table and figures for "Spatial regulation of coordinated excitatory and inhibitory synaptic plasticity at dendritic synapses"

| REAGENT or RESOURCE | SOURCE | IDENTIFIER |
|---|---|---|
| Antibodies | | |
| Anti-vGAT-Oyster 550 | Synaptic System | 131103C3, RRID:AB_887867 |
| Rat monoclonal anti-HA | Roche | 1186742300, RRID:AB_10094468 |
| Qdot® 655 F(ab')2-Goat anti-Rat IgG (H+L) | Thermo Fisher | Q-11621MP, RRID:AB_2556477 |
| Bacterial and Virus Strains | | |
| Biological Samples | | |
| Chemicals, Peptides, and Recombinant Proteins | | |
| BAPTA | Sigma | 85233-19-8 |
| L-NAME | Sigma | N5751 |
| Nifedipine | Sigma | N-7634 |
| Bicuculline | Sigma | 14343 |
| MDL28170 | Sigma | M-6690 |
| KN-93 | Millipore Merck | 422708 |
| KN-92 | Millipore Merck | 422709 |
| APV | Tocris | 0105/50 |
| CNQX | Tocris | 1045/10 |
| ω-conotoxin MVIIC | Tocris | 1084/100U |
| ω-conotoxin GVIA | Tocris | 1085/250U |
| DPNI-caged-GABA | Tocris | 2991/10 |
| MNI-caged-L-glutamate | Tocris | 1490/10 |
| Rhod-2 tripotassium salt | AAT Bioquest | 21067 |
| Critical Commercial Assays | | |
| Effectene Transfection Reagent | Qiagen | 301427 |
| Deposited Data | | |

| | | |
|---|---|---|
| **Experimental Models: Cell Lines** | | |
| **Experimental Models: Organisms/Strains** | | |
| Ai9 (B6.Cg-*Gt(ROSA)26Sortm9(CAGtdTomato) Hze*/J | Jackson Laboratory, USA | JAX:007909, RRID:IMSR_JAX:007909 |
| PVCRE (B6;129P2-Pvalbtm1(cre) Arbr/J | Jackson Laboratory, USA | JAX:017320, RRID:IMSR_JAX:017320 |
| Parvalbumin-tdTomato (PV-tdTomato) | This paper | |
| **Oligonucleotides** | | |
| **Recombinant DNA** | | |
| pEGFP-N1 | Clontech | Cat# 632162 |
| pcDNA3 Homer1c::DsRed | Petrini et al., 2009 | N/A |
| pcDNA3 Homer1c::GFP | Petrini et al., 2009 | N/A |
| FingR-Gephyrin-GFP | Gross et al., 2013 | N/A |
| EGFP-Gephyrin | Zita et al., 2007 | N/A |
| Hemagglutinin (HA)-tagged α1 GABAA receptor | | |
| **Software and Algorithms** | | |
| Metamorph 7.8 | Molecular Devices | http://www.moleculardevices.com/Products/Software/Meta-Imaging-Series/MetaMorph.html RRID: SCR_002368 |
| Clampex 10.6 | Molecular Devices | http://www.moleculardevices.com/products/software/pclamp.html RRID:SCR_011323 |
| Clampfit 10.7 | Molecular Devices | http://www.moleculardevices.com/products/software/pclamp.html RRID:SCR_011323 |
| MATLAB | Mathworks | http://www.mathworks.com/products/matlab/ RRID:SCR_001622 |

| | | |
|---|---|---|
| GraphPad Prism 6 | GraphPad | http://www.graphpad.com/<br>RRID: SCR_002798 |
| Custom program written for MATLAB to reconnect QD trajectories | Petrini et al., 2014; from<br>D Choquet and L Cognet | N/A |
| Custom program for SPT quantifications | Petrini et al., 2014; from<br>D Choquet and A Serge | N/A |
| **Other** | | |

### SUPPLEMENTARY FIGURES

#### Figure S1

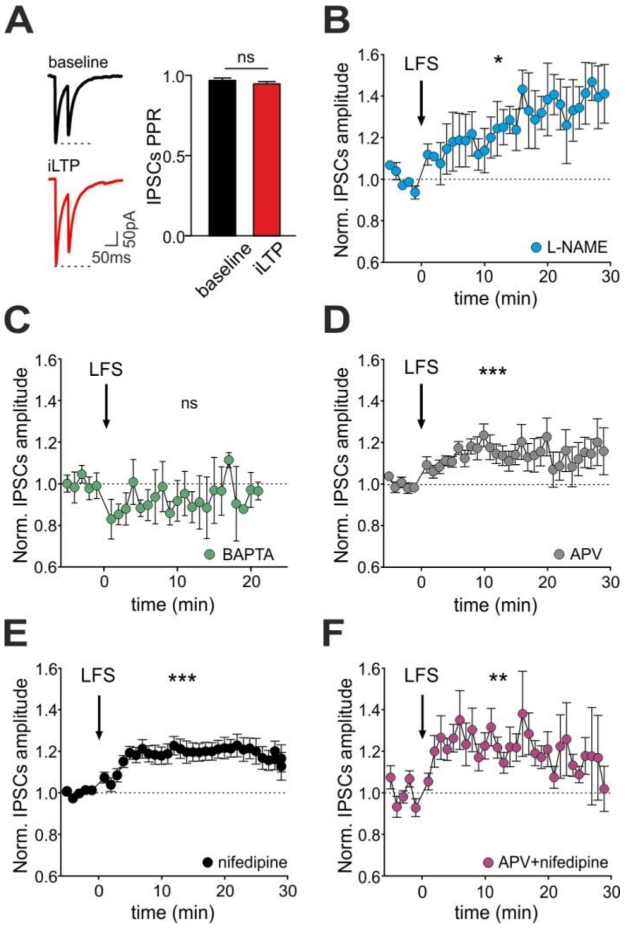

**Figure S1 (related to Figure 1): Postsynaptic mechanism and $Ca^{2+}$ dependence of LFS-induced iLTP**

**A-B.** Unlikely presynaptic mechanisms of iLTP. **A.** Left: Representative IPSC paired pulses traces recorded before (baseline) and after (iLTP) the delivery of the LFS. Right: Quantification of the paired pulse ratio (PPR) (n = 25; p = 0.14, paired Student's t-test). **B.** The nitric oxide synthase blocker L-NAME does not prevent LFS-induced iLTP (n = 5, $F_{33,136}$= 1.6, p = 0.03; one-way ANOVA followed by Turkey's multiple comparison test). **C-E.** Time course of relative IPSC amplitude increase before and after the delivery of the LFS protocol (arrow), in the presence of the fast $Ca^{2+}$ chelator BAPTA (**C**; n = 4, $F_{25,69}$ = 0.4, p = 0.99), APV (**D**; n = 11, $F_{33,241}$ = 2.2, p < 0.001), nifedipine (**E**; n = 21, $F_{33,640}$ = 3.6, p < 0.001), and APV + nifedipine (**F**; n = 6, $F_{33,162}$ = 2.1, p = 0.002). One-way ANOVA followed by Turkey's multiple comparison test. Values are expressed as mean ± SEM. *p < 0.05, **p < 0.01, ****p < 0.001, ns = not significant.

### Figure S2

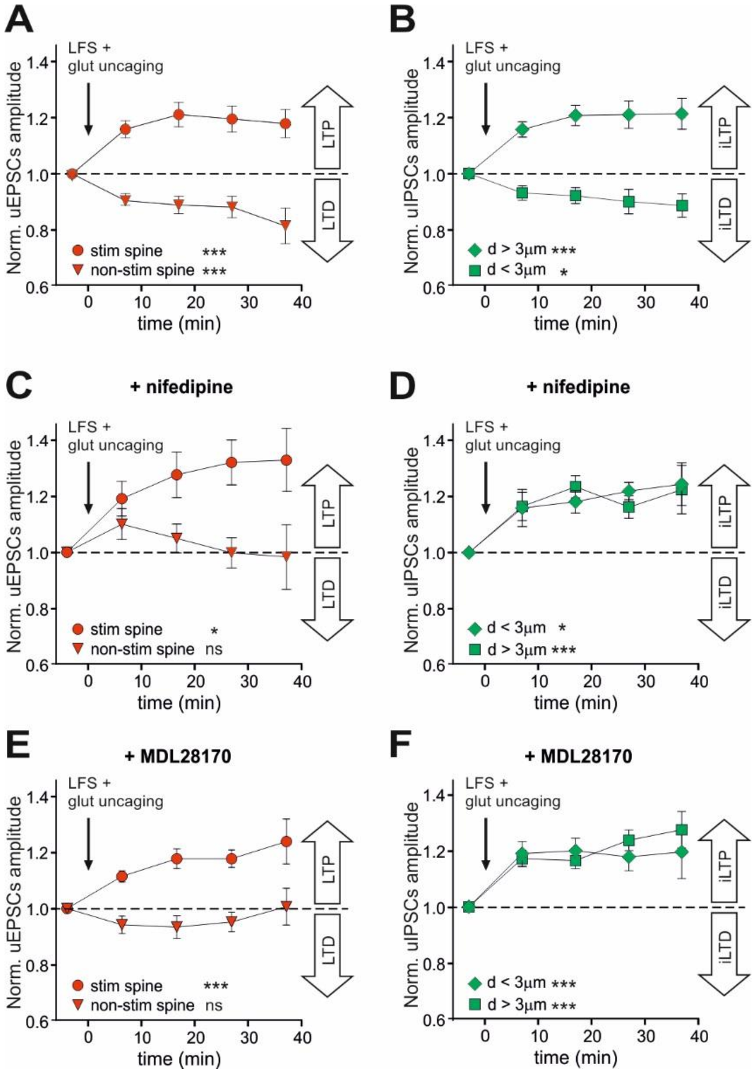

**Figure S2 (related to Figure 4): Spatial coordination of the plasticity of excitatory and inhibitory synapses upon single spine LTP**

**A.** After the "LFS+glut uncaging" protocol, the stimulated spine is selectively potentiated (circle, n = 7-20 synapses from 20 neurons, $F_{4,61} = 9.3$, $p < 0.001$) and the non-photostimulated ("non-stim") spines (putatively exposed only to LFS) are depressed (triangle, n = 6-16 from 20 neurons, $F_{4,51} = 6.3$, $p < 0.001$). All the statistical comparison shown here are performed with one-way ANOVA followed by Dunnett's post-test. **B.** After the "LFS+glut uncaging protocol", GABAergic synapses located at d > 3 µm from the stimulated spine are potentiated (diamond, n = 7-41 synapses from 20 neurons, $F_{4,127} = 11.4$, $p < 0.001$) and those located at d < 3 µm are depressed (square, n = 11-30 20 neurons, $F_{4,103} = 3.0$, $p = 0.02$). **C.** Same as in A in presence of nifedipine. Stimulated spine, n = 4-7 synapses from 7 neurons, $F_{4,26} = 3.9$, $p = 0.01$; non-photostimulated spine, n = 3-6 synapses from 7 neurons, $F_{4,20} = 0.8$, $p = 0.51$. **D.** Same as in B, in presence of nifedipine. d < 3 µm, n = 3-9 synapses from 7 neurons, $F_{4,27} = 4.0$, $p = 0.01$; d > 3 µm, n = 4-14 synapses from 7 neurons, $F_{4,45} = 6.1$, $p < 0.001$. **E.** Same as in A in presence of MDL28170. Stimulated spine, n = 3-24 synapses from

24 neurons, $F_{4,78}$ = 11.9, p <0.001; non-photostimulated spine, n = 3-25 synapses from 24 neurons, $F_{4,70}$ = 1.2, p = 0.31. **F.** Same as in B, in presence of MDL28170. d < 3 µm, n = 5-24 synapses from 24 neurons, $F_{4,71}$ = 7.1, p <0.001; d > 3, n = 6-68 synapses from 24 neurons, $F_{4,174}$ = 20.0, p <0.001. Values are expressed as mean ± SEM. *p < 0.05, ***p < 0.001, ns = not significant.

### Figure S3

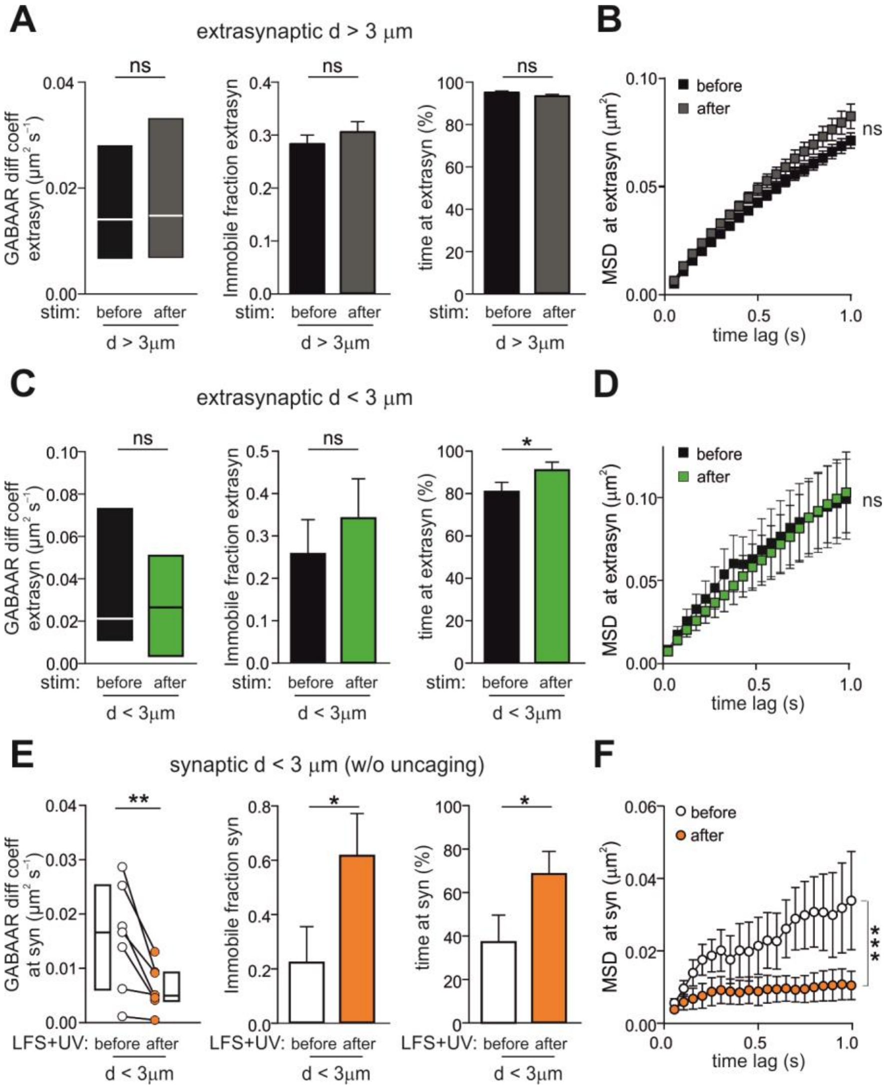

**Figure S3 (related to Figure 7): Supplementary data on the modulation of GABAAR lateral mobility upon single spine LTP**

**A-B.** Characterization of the lateral mobility of extrasynaptic GABAARs located at d > 3 µm from the potentiated spine, before (black) and after (grey) the single spine LTP protocol. **A**. Left: Median diffusion coefficient and interquartile range (IQR; n = 526-620 trajectories from 22 neurons; p = 0.63, Mann-Whitney test). Middle: Immobile fraction (n = 526-620 trajectories from 22 neurons; p = 0.40, Mann-Whitney test). Right: Percentage of time spent by GABAA receptors in the extrasynaptic compartment (n = 526-638 trajectories; $p = 0.16$, Mann-Whitney test). **B.** MSD versus time plot (n= 526-617 from 22 neurons; ns, ordinary two-way ANOVA followed by Bonferroni's post hoc test). **C-D.** Characterization of the lateral mobility of extrasynaptic GABAARs located at $d < 3$ µm from the stimulated spine, before (black) and after (green) the single spine LTP protocol. **C**. Left: Paired median diffusion coefficient ($n = 25$ trajectories from 14 neurons; $p = 0.34$, paired Wilcoxon test). Middle: Paired IF ($n = 25$ trajectories from 14 neurons; $p = 0.24$, paired Wilcoxon test). Right: Paired values of percentage time spent by GABAA receptors in the extrasynaptic compartments at $d < 3$ µm from the stimulated spine ($n = 25$ trajectories from 14 neurons; $p = 0.01$, paired Wilcoxon test). **D.** MSD versus time plot of paired extrasynaptic GABAA receptors close to the potentiated spine ($d < 3$ µm), $n = 18$ from 14 neurons, ns, RM two-way ANOVA followed by Bonferroni's post hoc test. **E-F**. Same as in C-D, except for the uncaging. Please note that in this set of experiments the stimulating protocol was LFS + 4Hz UV-light pulses train on a spine (ctrl spine) in absence of MNI-glutamate. Only synaptic GABAAR trajectories localized in the range of 3µm from the ctrl spine were considered. **E**. Left: Paired median diffusion coefficient ($n = 7$ from 4 neurons; $p = 0.01$, paired Wilcoxon test). Middle: Paired IF ($n = 7$ from 4 neurons; $p = 0.03$, paired Wilcoxon test). Right: Paired values of percentage of time spent by GABAA receptors at synapses close to the control spine ($n = 7$ from 4 neurons; $p = 0.01$, paired Wilcoxon test). **F.** Paired MSD values of synaptic GABAA receptors close to the control spine ($d < 3$ µm; $n = 4$ from 4 neurons, $p < 0.001$, RM two-way ANOVA). Unless stated otherwise, values are expressed as mean ± SEM. $*p < 0.05$, $**p < 0.01$, ns = not significant.
